# Self-regulatory function of bacterial small heat shock protein IbpA through mRNA binding is conferred by a conserved arginine

**DOI:** 10.1101/2023.04.26.538363

**Authors:** Yajie Cheng, Tsukumi Miwa, Hideki Taguchi

## Abstract

Bacterial small heat shock proteins, IbpA and IbpB, co-aggregate with denatured proteins and recruit other chaperones for the processing of aggregates, thereby assisting in their refolding. In addition, as a recently revealed uncommon feature, *Escherichia coli* IbpA self-represses its own translation through interaction with the 5’ untranslated region (UTR) of the *ibpA* mRNA, enabling IbpA to act as a mediator of negative feedback regulation. Although IbpA also suppresses the expression of IbpB, IbpB does not have the self-repression activity despite the two Ibps being highly homologous. This study demonstrates that the self-repression function of IbpA is conserved in other bacterial IbpAs. Moreover, a cationic residue-rich region in the α-crystallin domain (ACD) of IbpA, which is not conserved in IbpB, is critical for the self-suppression activity. Notably, arginine 93 (R93) located within the ACD is an essential residue that cannot be replaced by the other 19 amino acids, including lysine. IbpA-R93 mutants completely lost the interaction with the 5’ UTR of the *ibpA* mRNA but retained almost all of the chaperone activity to sequester denatured proteins. Taken together, the conserved Arg93-mediated translational control of IbpA through RNA binding would be beneficial for a rapid and massive supply of the chaperone on demand.

## Introduction

Small heat shock proteins (sHsps) are characterized by their low molecular weights (12∼43 kD) in the subunits and conserved α-crystallin domains (ACDs) franked by disordered N-terminal domains (NTDs) and C-terminal domains (CTDs) (1–4). As molecular chaperones, sHsps are widely conserved in all kingdoms of life to protect cellular protein homeostasis (proteostasis) by binding with and then sequestering misfolded proteins in an ATP-independent manner, which is termed as a ‘sequestrase’ activity (4). Other chaperones, such as Hsp70 family, are required for later refolding or degradation to disassociate co-aggregates between sHsps and substrate proteins since sHsps are not refolding-active (1, 2, 4).

sHsps assemble into dimers and large oligomers (1–4). A basic scaffold of sHsps is formed through the ACD dimerization, with an ACD interacting with a partnering ACD to form homo- or hetero-dimers (1–4). The higher-order oligomers are then regulated by intrinsically disordered NTDs and CTDs (1–4). The CTD tails harbor a short yet highly-conserved functional sequence motif, the IXI/V motif, which plays a crucial role in the oligomerization and chaperone activity of sHsps (1–5). In bacterial sHSPs, the IXI/V motif interacts with a β4/β8 groove of the neighboring ACD to assemble two dimers (2, 4, 5). Mutations in the IXI/V motif, such as IXI-to-AEA, usually cause defects in the oligomerization ability and chaperone activity (2, 6). Notably, sHsp subunit associations are relatively weak, and oligomeric sHsps are primarily in a dimer-based and dynamic subunit exchange (4, 5). This property enables sHsps to cope with environmental changes such as heat stress, pH, and oxidative stress (4, 5).

sHsps can rapidly respond to environmental stresses, as evidenced by the significant upregulation of expression (7). Bacterial sHsps are upregulated by a sigma factor σ^32^, at the transcriptional level upon heat shock (8). Moreover, in α- and γ-proteobacteria, sHsps are translationally regulated by a thermo-sensitive mRNA structure (RNA thermometer, RNAT) in the 5’-untranslated region (5’ UTR), which contains the element of heat shock gene expression repression (9, 10). The elements, ranging in length from 60 to over 100 nucleotides, typically consist of 2 to 4 stem-loops, which play a critical role in the modulation of sHsp translation (11).

The number of sHsp members in organisms varies from one or two in prokaryotes to even over ten in eukaryotes (4). In *Escherichia coli*, two sHsps, inclusion body-associated protein A (IbpA_Ec_) and B (IbpB_Ec_), encoded in the *ibpAB* operon, share high-similarity amino acid sequences (∼ 50% identity, Fig. 1A) but are specialized in different functions during the anti-aggregation process (12). IbpA_Ec_ and IbpB_Ec_ interact with misfolded client proteins to form co-aggregates (so-called holdase activity) but are not involved in the subsequent refolding and degradation processes (13). The co-aggregates are recognized by other chaperones such as DnaK/DnaJ and ClpB for refolding/degradation, at which point IbpAB are released from the substrates (14). IbpA_Ec_ is more efficient for associating with client proteins to form small co-aggregates, while IbpB_Ec_, forming functional complexes with IbpA_Ec_, is more competent to assist the disassembly of sHsps from the co-aggregates (12, 15). IbpA_Ec_ and IbpB_Ec_ can form hetero-dimers and hetero-oligomers (16). In the absence of IbpB_Ec_, IbpA_Ec_ tends to form fibril-like structures both in vivo and in vitro, which might be mediated by NTD and CTD (6, 17). The fibrillation of IbpA_Ec_ can be blocked by its substrate proteins as well as IbpB_Ec_ (17).

**Fig. 1.**
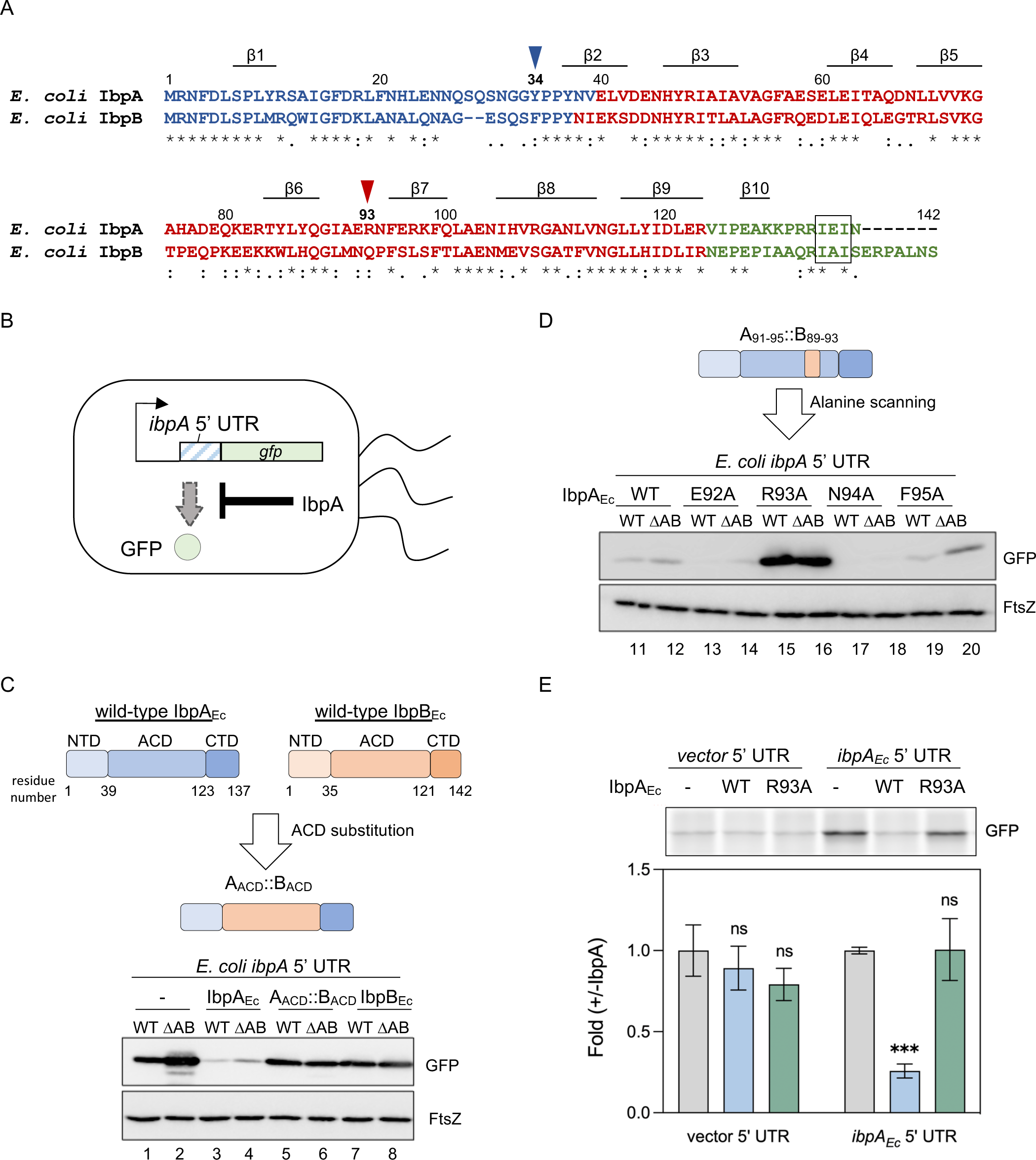
Identification of a crucial residue, Arg93, in *E. coli* IbpA (IbpA_Ec_) for discriminating between IbpA_Ec_ and IbpB_Ec_ in IbpA-mediated translation suppression function. (*A*) Alignment of *E. coli* IbpA (IbpA_Ec_) and IbpB (IbpB_Ec_). NTDs, ACDs, and CTDs are colored blue, red, and green, respectively. IbpA_Ec_ tyrosine 34 (Y34) and arginine 93 (R93) are indicated by blue and red arrowheads, respectively. The IXI/V motif, located in CTDs, are marked by a black box. Locations of β strands, based on an AlphaFold2-predicted IbpA_Ec_ monomer structure, are also shown. (*B*) IbpA-mediated translation suppression using a GFP reporter assay. The translation of GFP reporter from *gfp* mRNA harboring the 5’ UTR of *ibpA_Ec_* in *E. coli* was inhibited by excess amounts of IbpA. (*C*, *D*) Systematic chimera and mutagenesis to identify a critical residue in IbpA_Ec_ to differentiate between IbpA_Ec_ (*blue*) and IbpB_Ec_ (*orange*) using the GFP reporter assay. *Upper*: Schematic of an IbpA_Ec_/IbpB_Ec_ chimera *(C)* and subsequent single-residue IbpA_Ec_ mutations (*D*). *Lower*: Western blotting analysis evaluating the effects of the ACD substitution *(C)* and individual alanine substitution among IbpA_Ec_ residues 92∼95 (*D*) on the reporter GFP translation level in the *E. coli* BW25113 strains (WT: BW25113 wild-type strain; ΔAB: *ibpAB* operon-deleted BW25114 strain). The expression of FtsZ was used as a control of the constitutive expression level. Anti-GFP and anti-FtsZ antibodies were used for the detection. Note: other chimera and mutation analyses are shown in Fig. S1 and S2. (*E*) Cell-free translation in the absence or presence of purified IbpA_Ec_-WT or the R93A mutant. The *gfp* reporter mRNAs carrying a vector or the *ibpA_Ec_* 5’ UTR were translated in the *E. coli* reconstituted cell-free translation system (PURE system). *Upper*: the fluorescence intensity of the translated GFP; *lower*: the fold of quantified GFP fluorescence level in the presence of IbpA_Ec_ (WT or R93A) compared to that in the absence of IbpA_Ec_ (-). The data represent the means (±S.D.) of three independent experiments and were analyzed by one-way ANOVA within the comparison with each -IbpA group (*ns*: non-significant; \**p*<0.0332; \*\**p*<0.0021; \*\*\**p*<0.0002; \*\*\*\**p*<0.0001, wherever shown).

Although IbpA_Ec_ and IbpB_Ec_ are highly similar in sequence and closely collaborate in both structures and chaperone activity, IbpA_Ec_ possesses a unique post-transcriptional regulation mechanism that IbpB_Ec_ lacks (18). IbpA_Ec_ is up-regulated at the posttranscriptional level even without heat stress by the overexpression of aggregation-prone proteins (18). Further analysis revealed that IbpA_Ec_ directly interacts with the 5’ UTR of *ibpA* (and *ibpB*) mRNA and inhibits the translation of *ibpA* (and *ibpB*). The IbpA-mediated translation suppression is relieved by aggregation-prone client proteins that recruit IbpA_Ec_, thereby reducing the amount of IbpA_Ec_ involved in self-suppression (18). The nonconventional function of IbpA_Ec_ as an aggregation-sensor tightly suppresses IbpA_Ec_ expression under aggregation-free conditions but enables cells to rapidly up-regulate the IbpA_Ec_ levels upon acute aggregation stress, such as heat shock (10, 18).

Elucidating the molecular mechanism underlying IbpA_Ec_ self-suppression on translation is of great interest, but many questions remain. For example, why does IbpB_Ec_ lack self-regulation activity? Is this translational repression by IbpA conserved in other bacterial IbpAs? Here, we revealed the conservation of the IbpA-mediated self-regulation in other bacterial IbpAs and found that a highly-conserved residue, Arg93 (R93), located within the IbpA ACD and absent in IbpB, is irreplaceably important for discriminating between IbpA and IbpB in the translation suppression function.

## Results

### The residue Arg93, located within the α-crystallin domain (ACD), plays a crucial role in the self-regulation of IbpA_Ec_

To explore the region responsible for the notable difference between IbpA_Ec_ and IbpB_Ec_, we conducted a reporter assay to assess the translation regulation activity of IbpA_Ec_ (18). In brief, the translation of a *gfp* reporter, containing the *ibpA_Ec_*5’ UTR sequence, is suppressed and increased in *E. coli* cells with overexpressed and deleted IbpA_Ec_, respectively (18) (Fig. 1B).

We overexpressed chimeric IbpABs that included ACD, NTD, and CTD domain substitutions in both *E. coli* wild-type (WT) and the *ibpAB* operon-deleted (ΔAB) strains, along with reporter plasmids (Fig. 1C, Fig. S1). The expression of the GFP reporter was considerably increased in the absence of endogenous IbpA_Ec_ (Fig. 1C, lane 2), Conversely, its expression was markedly reduced due to the overexpression of exogenous IbpA_Ec_-WT (Fig. 1C, lanes 3 and 4), confirming the suppressive effect of IbpA_Ec_ on the reporter translation, as shown previously (18). The chimeric IbpABs with substitutions in both NTD and CTD had no significant effects on self-suppression (Fig. S1). In contrast, the A_ACD_::B_ACD_ chimera, in which the IbpA_Ec_ ACD was replaced with that of IbpB_Ec_, significantly abolished the translation suppression (Fig.1C, lanes 5 and 6), indicating that the IbpA_Ec_ ACD includes a critical region for the self-regulation activity of IbpA_Ec_. Following a series of mutation experiments in ACD (Fig. S2), we narrowed the critical region down to IbpA_Ec_ residues from 92 to 95 and then individually mutated these residues with alanine. Among these mutants, mutation of Arg93 to Ala (IbpA_Ec_-R93A) greatly increased the translation level of the reporter *gfp*, compared to the WT and the other alanine mutants (Fig. 1D, lanes 15, 16), indicating that the point mutation R93A is sufficient to lose the translation suppression activity.

Next, we evaluated the effect of IbpA_Ec_-R93A using a chaperone-free, reconstituted cell-free translation system (PURE system) (19), as previously used for the IbpA_Ec_-mediated translation suppression (18). Purified IbpA_Ec_-WT suppressed the translation of the *gfp* reporter in an *ibpA_Ec_*5’ UTR-dependent manner (18), whereas purified IbpA_Ec_-R93A had no suppressive effect on the translation level (Fig. 1E), providing direct evidence that the IbpA_Ec_-R93A lost the ability to suppress its own translation.

### Conservation of self-regulation activity and the significance of R93 in other bacterial IbpAs

To investigate the conservation of translational repression activity and the importance of R93 in IbpA_Ec_ across other bacterial IbpAs, we selected two IbpAs from *Cedecea neteri* (IbpA_Cn_) and *Vibrio harveyi* (IbpA_Vh_) (15). Although the homology of these two IbpAs differs from IbpA_Ec_ (15), R93 is conserved in all IbpAs (Fig. 2A). We overexpressed IbpA_Cn_ and IbpA_Vh_ or the corresponding R93 mutants (IbpA_Cn_-R93A and IbpA_Vh_-R94A) in *E. coli* with GFP-reporter plasmids harboring the *ibpA_Cn_* and *ibpA_Vh_*5’ UTRs, respectively, which conform to the structural properties of RNAT (Fig. S3, Table S1) (11). We found that the overexpression of IbpA_Cn_ and IbpA_Vh_ in *E. coli* suppressed the expression of GFP, which was relieved by the corresponding R93 mutants (Fig. 2B). Furthermore, in the PURE system analysis, purified IbpA_Cn_ and IbpA_Vh_ WT suppressed the translation of the *gfp* reporter, whereas the corresponding R93 mutants had no suppression effect (Fig. 2C). Together, the results show the prevalence of the IbpA-mediated self-translation inhibition activity and the general importance of R93 in IbpAs.

**Fig. 2.**
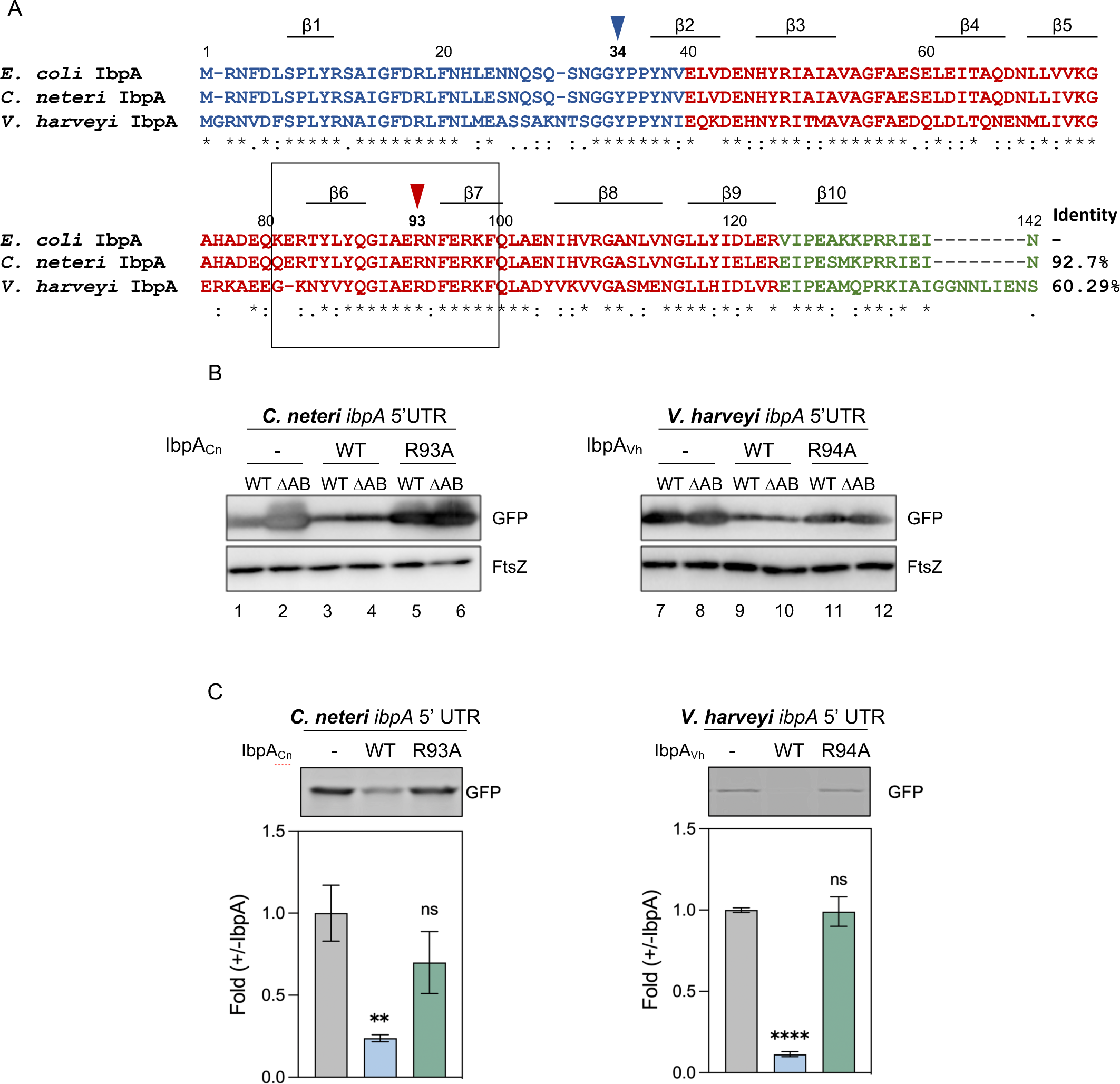
Conservation of self-translation regulation and the critical residue in other bacterial IbpAs. (*A*) Multiple sequence alignment of *E. coli* IbpA (IbpA_Ec_), *C. neteri* IbpA (IbpA_Cn_), and *V. harveyi* IbpA (IbpA_Vh_). NTDs, ACDs, and CTDs are represented in blue, red, and green, respectively. The amino acids Y34 and R93 are indicated by blue and red arrowheads, respectively. Arg/Lys-rich regions in IbpA ACDs are highlighted with a black box. IbpA_Cn_ and IbpA_Vh_ share 93% and 60% amino acid sequence identity with IbpA_Ec_, respectively. (*B*) Translation suppression activity of IbpA_Cn_ (*left*), IbpA_Vh_ (*right*) and their respective R93 mutants evaluated by the GFP reporter assay. The *gfp* mRNA harboring the 5’ UTRs of *ibpA_Cn_*(*left*) or *ibpA_Vh_* (*right*) were expressed in *E. coli* (WT: BW25113 wild-type strain; ΔAB: *ibpAB* operon-deleted BW25114 strain). The levels of GFP (reporter) and FtsZ (loading control) were detected by anti-GFP and anti-FtsZ antibodies, respectively. (*C*) Cell-free translation in the presence or absence of purified IbpA_Cn_ (*left*), IbpA_Vh_ (*right*), and their respective R93 mutants. The *gfp* reporter mRNAs carrying either the *ibpA_Cn_* or the *ibpA_Vh_* 5’ UTR were translated in the PURE system. The fluorescence intensity of translated GFP (*upper*) was quantified and normalized to the control group without IbpAs (-). The data represent the means (±S.D.) of three independent experiments and were analyzed by one-way ANOVA within the comparison with each -IbpA group (*ns*: non-significant; \**p*<0.0332; \*\**p*<0.0021; \*\*\**p*<0.0002; \*\*\*\**p*<0.0001, wherever shown).

### Arg93 in IbpA has irreplaceable importance on IbpA self-suppression

Previous studies on human sHsps have shown that a region in ACD enriched with positively-charged residues is associated with human congenital diseases (20, 21). For example, the R116C mutant of human HspB4 is linked to cataract disease, resulting in non-negligible changes in the structures, chaperone activity, and oligomerization trend (22), while the R116K mutant retains similar properties to HspB4 WT, indicating that the basic amino acid in that position is essential for maintaining HspB4 activities (23). We noticed that the positive-charge cluster in ACD is conserved in *E. coli*, *C. neteri,* and *V. harveyi* IbpAs (R83-K98 in *E. coli* IbpA, as shown in Fig. 2A and 3A) but not in IbpB. Consequently, we investigated the role of conserved basic residues in IbpA-mediated self-suppression. Reporter assays using alanine-substituted mutants (IbpA_Ec_-R83A, R97A, and K98A) demonstrated that R97A and K98A, but not R83A, lost IbpA-mediated translation suppression activity similar to R93A (Fig. 3B). We then mutated IbpA_Ec_-R93/R97/K98 to K/R to maintain the position’s positive charge. We found that R97K and K98R mutants maintained translation suppression activity comparable to IbpA_Ec_-WT (Fig. 3C, lanes 17-20), whereas R93K was repression-defective like R93A (Fig. 3C, lanes 15 and 16). The results demonstrate that the positive charges at positions 97 and 98 are sufficient to preserve IbpA suppression activity, but R93 is not replaceable with Lys. To further investigate the exclusive role of Arg in the 93^rd^ position in IbpA self-suppression, we mutated R93 in IbpA_Ec_ with all other amino acid residues individually. The reporter assay results showed that all other R93 mutants lost the suppression ability (Fig. S4), emphasizing the irreplaceable importance of R93 in IbpA self-regulation.

**Fig. 3.**
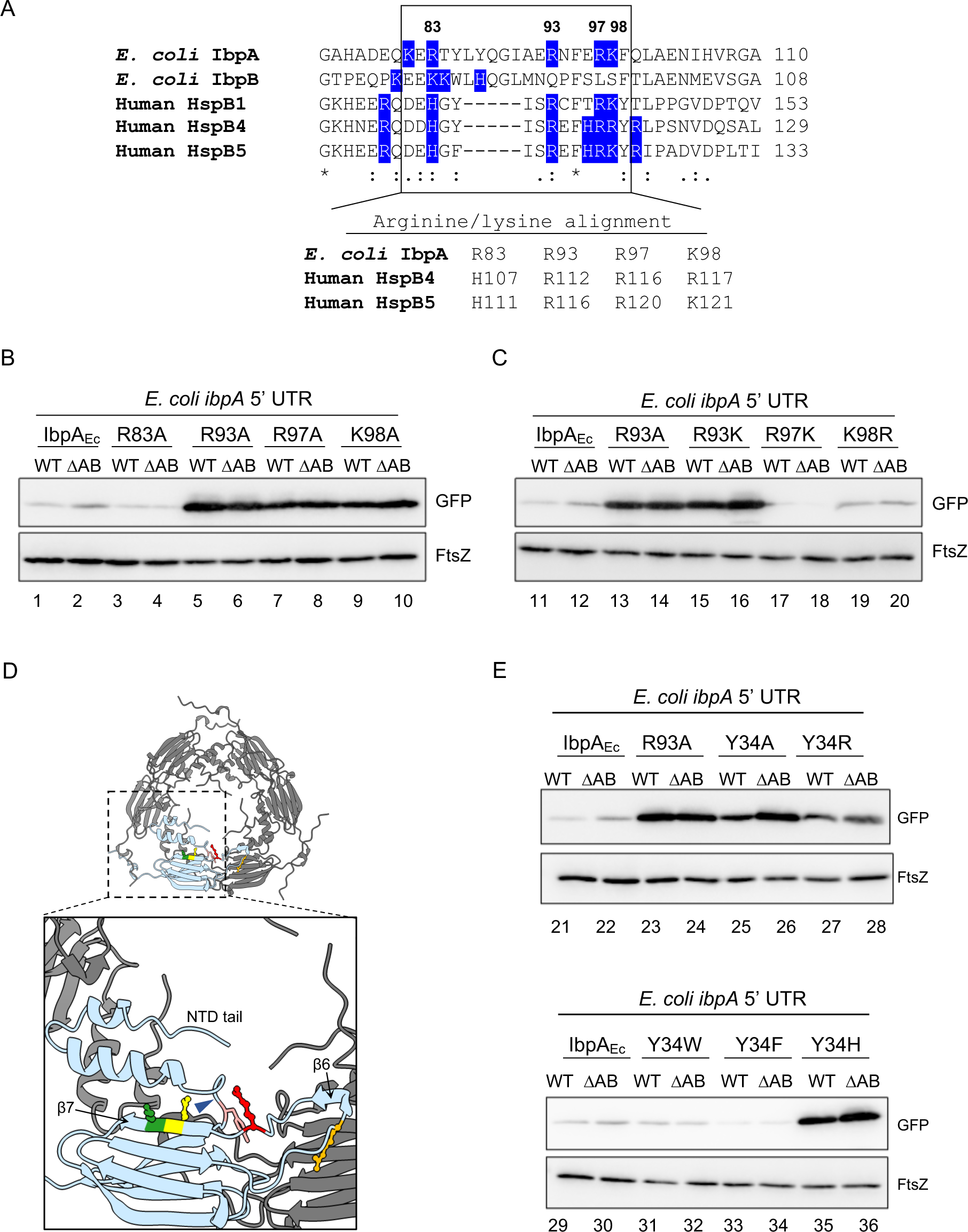
The effect of cationic amino acids near Arg93 on IbpA_Ec_-mediated self-regulation function. (*A*) Partial alignment of IbpA_Ec_, IbpB_Ec_, and human sHsps (HspB1, HspB4 and HspB5). The region enriched in cationic residues (R/K/H) is enclosed in a box. The conserved Arg/Lys residues in the boxed area are indicated below. *(B)* The effect of Ala mutants of R83, R93, R97, and K98 in IbpA_Ec_ on the GFP reporter harboring the *ibpA_Ec_* 5’ UTR. WT: *E. coli* BW25113 wild-type strain; ΔAB: *ibpAB* operon-deleted BW25114 strain. Anti-GFP and Anti-FtsZ were used to detect GFP and FtsZ levels. (*C*) Effect of Arg-to-Lys or Lys-to-Arg mutants of R93, R97, and K98 in IbpA_Ec_ on the GFP reporter. The experimental details are the same as in (B). (*D*) A hexamer structure of IbpA_Ec_-WT predicted by AlphaFold2 using MMseqs2 (33). The structure depicts one of the subunits in light blue for clarity. A zoom-in figure of the subunit is shown below, and the residues Y34, R83, R93, R97 and K98 are colored pink, orange, red, yellow, and green, respectively. Tyr34 in the NTD loop, which might interact with R93, is marked by a blue arrowhead. (*E*) The effect of IbpA_Ec_ Y34 mutants (Y34A, Y34R, Y34W, Y34F, and Y34H) on the GFP reporter was evaluated in *E. coli*, as described in (*B*).

What distinguishes R93 from R97/K98? The AlphaFold2-predicted structure of IbpA_Ec_ provides a possible explanation; while R93 is located in a flexible loop connecting β6 and β7 sheets, R97/K98 are positioned in the β7 sheet (Fig. 3D). Moreover, R93 resides in close proximity to a disordered NTD loop, implying a potential interaction with NTD. Similar interactions are suggested in the predicted structures of IbpA_Cn_ and IbpA_Vh_ (Fig. S5), indicating a common structural feature of R93. We then mutated Y34 in IbpA_Ec_, which is the closest residue to R93 in the predicted structure (Fig. 3D) and is highly conserved in IbpAs (Fig. 2A), to several amino acids. Notably, IbpA_Ec_-Y34A, Y34R, and Y34H mutants showed a loss of translation suppression activity, while Y34W and Y34F retained this activity (Fig. 3E), suggesting that aromatic residues at position 34 are crucial for the self-regulation function of IbpA. Taken together, these results suggest that the potential interplay between R93 and the NTD loop contributes to the self-suppression ability of IbpA.

### The self-regulation of IbpA is not solely dependent on its oligomer size

Given that the function of IbpA relies on its oligomeric structure, we investigated whether R93 mutation could induce conformational changes in IbpA_Ec_. Far-UV circular dichroism (CD) spectra of the purified IbpA_Ec_-R93A and IbpA_Ec_-WT were almost identical in the wavelength region that determines the secondary structure (Fig. S6A, B), but sucrose density gradient (SDG) centrifugation analysis revealed that IbpA_Ec_-R93A was mainly found at the bottom fraction compared to IbpA_Ec_-WT (Fig. 4A), indicating that IbpA_Ec_-R93A forms larger assemblies. Transmission electron microscopy (TEM) analysis showed that IbpA_Ec_-WT formed fibril-like structures (Fig. 4B), consistent with previous findings (17). IbpA_Ec_-R93A formed much longer fibrils than WT (Fig. 4B), in agreement with the tendency to form larger assemblies in the SDG analysis.

**Fig. 4.**
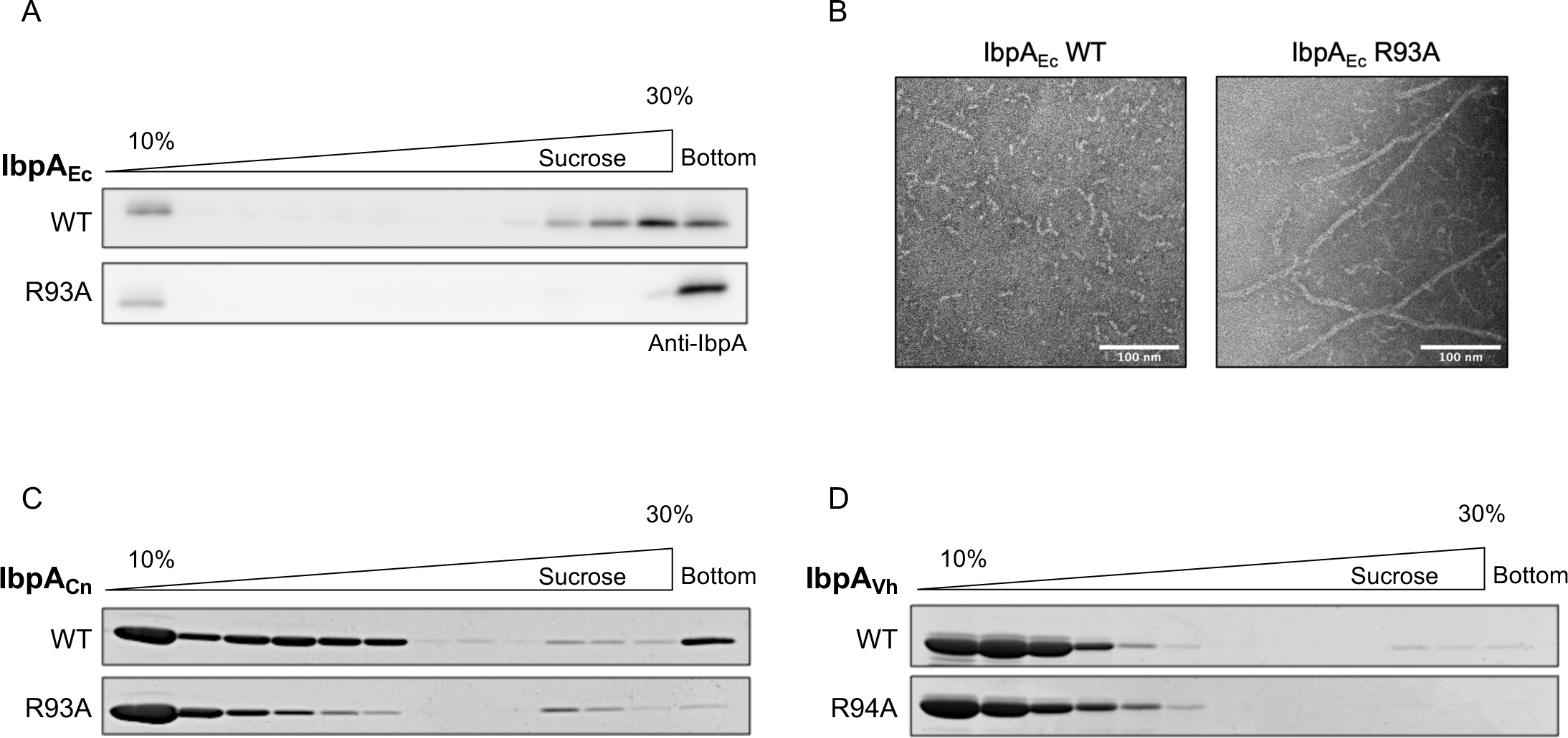
The self-regulation of IbpA is not solely dependent on oligomer size. (*A*) Oligomeric states of IbpA_Ec_ were analyzed using sucrose density gradient (SDG) centrifugation. Purified IbpA_Ec_-WT or R93A were applied to 10%-30% (w/v) sucrose gradient solutions, followed by SDS-PAGE and western blotting. The distribution of IbpA_Ec_ was probed with an anti-IbpA antibody. (*B*) Representative TEM images of purified IbpA_Ec_-WT (*left*) and R93A (*right*) are shown, with scale bars of 100 nm. (*C*, *D*) Oligomeric states of IbpA_Cn_ (*C*) and IbpA_Vh_ (*D*) (WT and their respective R93 mutants) were evaluated using SDG centrifugation and SDS-PAGE with CBB staining to visualize the IbpA distribution, as above.

We also examined the oligomeric states of IbpA_Cn_, IbpA_Vh,_ and their corresponding R93 mutants via SDG centrifugation. Unlike IbpA_Ec_, there was no significant difference in the oligomer distribution between WT and the mutants (Fig. 4C, D), suggesting that IbpA self-regulation does not solely rely on the oligomer size. The tendency of IbpA_Ec_-R93A to form larger assemblies was validated by the additional mutation analysis. In addition to the R93A mutation, we replaced a motif for oligomerization, IEI (IXI/V motif in IbpA_Ec_), in CTD with AEA. The IbpA-AEA mutant is known to disable higher-order oligomer formation (6), and SDG analysis confirmed the absence of oligomers in the IbpA_Ec_-AEA mutant. In contrast, the IbpA_Ec_-R93A/AEA still formed oligomers of various sizes (Fig. S7A). The in vivo reporter and PURE system analyses revealed that the IbpA-R93A/AEA was inactive in translation repression (Fig. S7B and C), supporting the idea that oligomer sizes are not the sole determinant for IbpA-mediated translation suppression activity.

### IbpA-R93A is impaired in the interaction with the *ibpA* 5’ UTR mRNA

We investigated the role of R93 in chaperone activity and mRNA interaction. Under conditions where all heated luciferases sedimented to the bottom in the SDG centrifugation, IbpA_Ec_-WT formed oligomers with heat-denatured luciferase in middle fractions (Fig. 5A). IbpA_Ec_-R93A also formed oligomers with luciferase in the middle fractions, although less efficiently than IbpA_Ec_-WT, indicating that the mutation of R93 partially impairs the chaperone function. We also evaluated the chaperone activity of IbpA_Cn_-R93A and IbpA_Vh_-R94A and found that they exhibited client-binding activity indistinguishable from the corresponding WT (Fig. 5A). These results suggest that the R93 mutation may weaken the chaperone activity but is not critical.

**Fig. 5.**
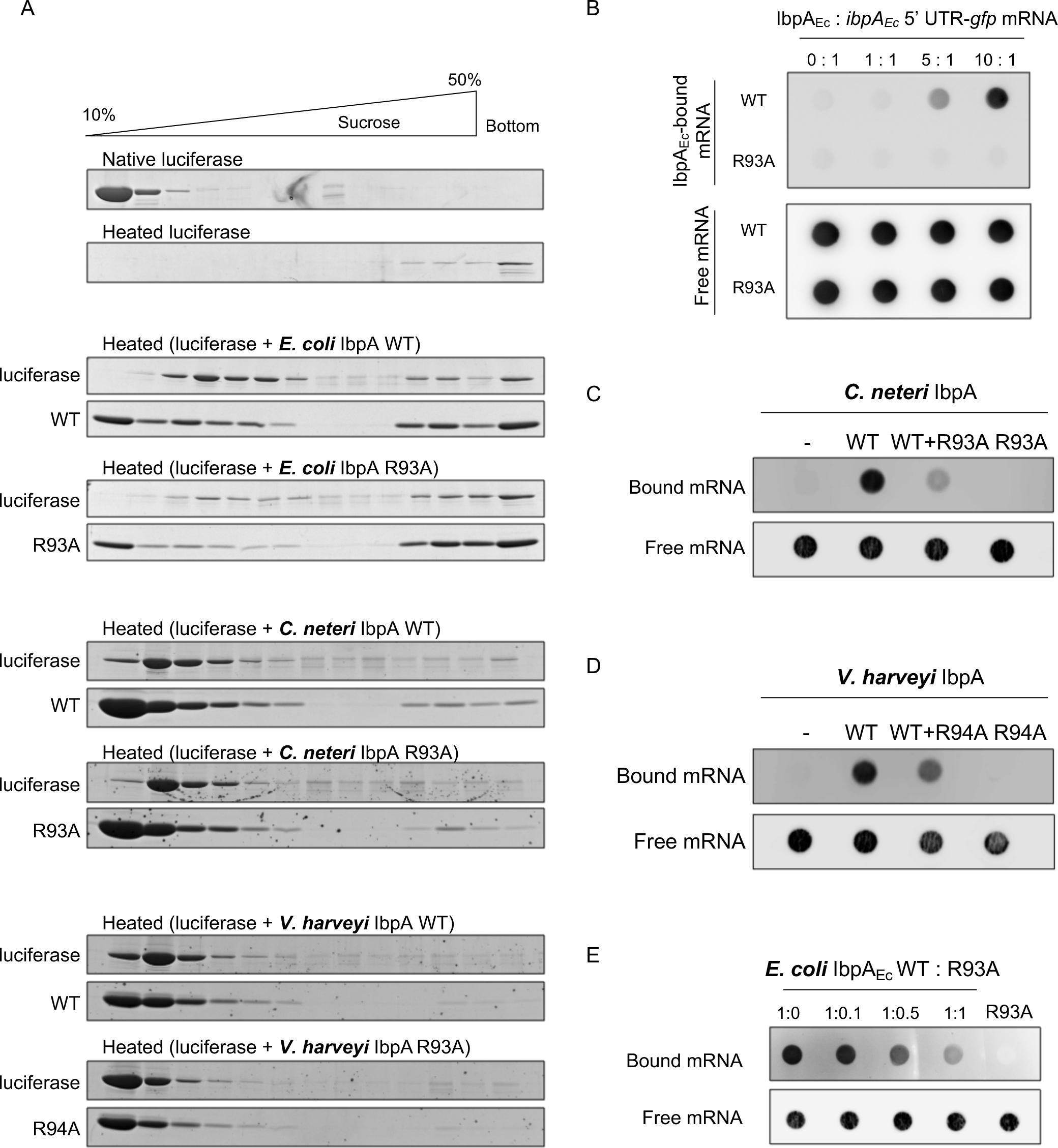
Interaction between IbpA-R93A mutants and denatured proteins or *ibpA* **5’ UTR mRNAs.** (*A*) The interaction between IbpAs and denatured luciferase was analyzed using SDG centrifugation. Luciferase was thermally denatured in the presence or absence of IbpA-WTs or respective R93 mutants, and then the mixtures were applied to 10%-50% (w/v) sucrose gradient solutions, followed by SDS-PAGE and CBB staining. (*B*) The interaction between IbpA_Ec_ and biotin-labeled *ibpA_Ec_*5’ UTR-*gfp* mRNA was evaluated using a filter binding assay. The mRNA and IbpA mixture was applied through double membranes, consisting of a nitrocellulose membrane (*upper*) and a positively-charged nylon membrane (*lower*), to capture IbpA-bound mRNA and free mRNA, respectively. mRNA intensity was detected by streptavidin-HRP. (*C*, *D*) The interaction of IbpA_Cn_-*ibpA_Cn_* 5’ UTR mRNA (*C*) and IbpA_Vh_-*ibpA_Vh_* 5’ UTR mRNA (*D*) was evaluated by the filter binding assay as described above. The applied amounts of the corresponding IbpA-R93 mutants were the same as the respective WTs. mRNA intensity was detected by the streptavidin Alexa Fluor 647 conjugate. (*E*) The filter binding assay was used to examine the interaction between IbpA_Ec_-WT and *ibpA* 5’ UTR-*gfp* mRNA in the presence of varying amounts of IbpA_Ec_-R93A. mRNA intensity was detected by streptavidin Alexa Fluor 647 conjugate.

Next, we tested the effect of R93 on the interaction with the *ibpA* 5’ UTR mRNA using a filter binding assay. When the mixture of proteins and RNAs is passed through a positively-charged nylon membrane covered with a nitrocellulose membrane, the protein-bound RNAs and free RNAs are trapped in the nylon and nitrocellulose membranes, respectively (24). The filter assay confirmed the interaction between purified IbpA_Ec_-WT and a biotin-labeled mRNA including the *ibpA* 5’ UTR in a dose-dependent manner, while there was no interaction of IbpA_Ec_-R93A with the mRNA (Fig. 5B). Combined with the results on the complete loss of the mRNA binding in IbpA_Cn_-R93A and IbpA_Vh_-R94A (Fig. 5C, D), we conclude that the R93 mutation loses the ability to bind the mRNA.

Furthermore, the addition of IbpA-R93A mutants weakened the mRNA binding ability of IbpA-WTs (Fig. 5C-E), suggesting the formation of a hetero-oligomer between IbpA-WTs and the R93A mutants causes the impaired interaction of IbpA-WTs with the mRNAs.

### IbpB enhances the suppression ability of IbpA-WT but has no effect on IbpA-R93A

In *E. coli*, IbpA_Ec_ and IbpB_Ec_ prefer the formation of hetero-species over homo-species (16) to mitigate aggregation-induced stress (4, 6). Subsequently, we investigated the impact of IbpB_Ec_-WT on IbpAB hetero-oligomer formation and IbpA-mediated translation suppression. SDG analysis showed that both IbpA_Ec_-WT and R93A formed hetero-complexes with IbpB_Ec_, which were smaller than corresponding homo-oligomers, particularly IbpA_Ec_-R93A (Fig. 6A). TEM analysis corroborated this trend; The addition of IbpB_Ec_ hindered the formation of fibril-like structures of IbpA_Ec_-WT, as demonstrated previously (17), and resulted in a shorter fibril formation in IbpA_Ec_-R93A (Fig. 6B).

**Fig. 6.**
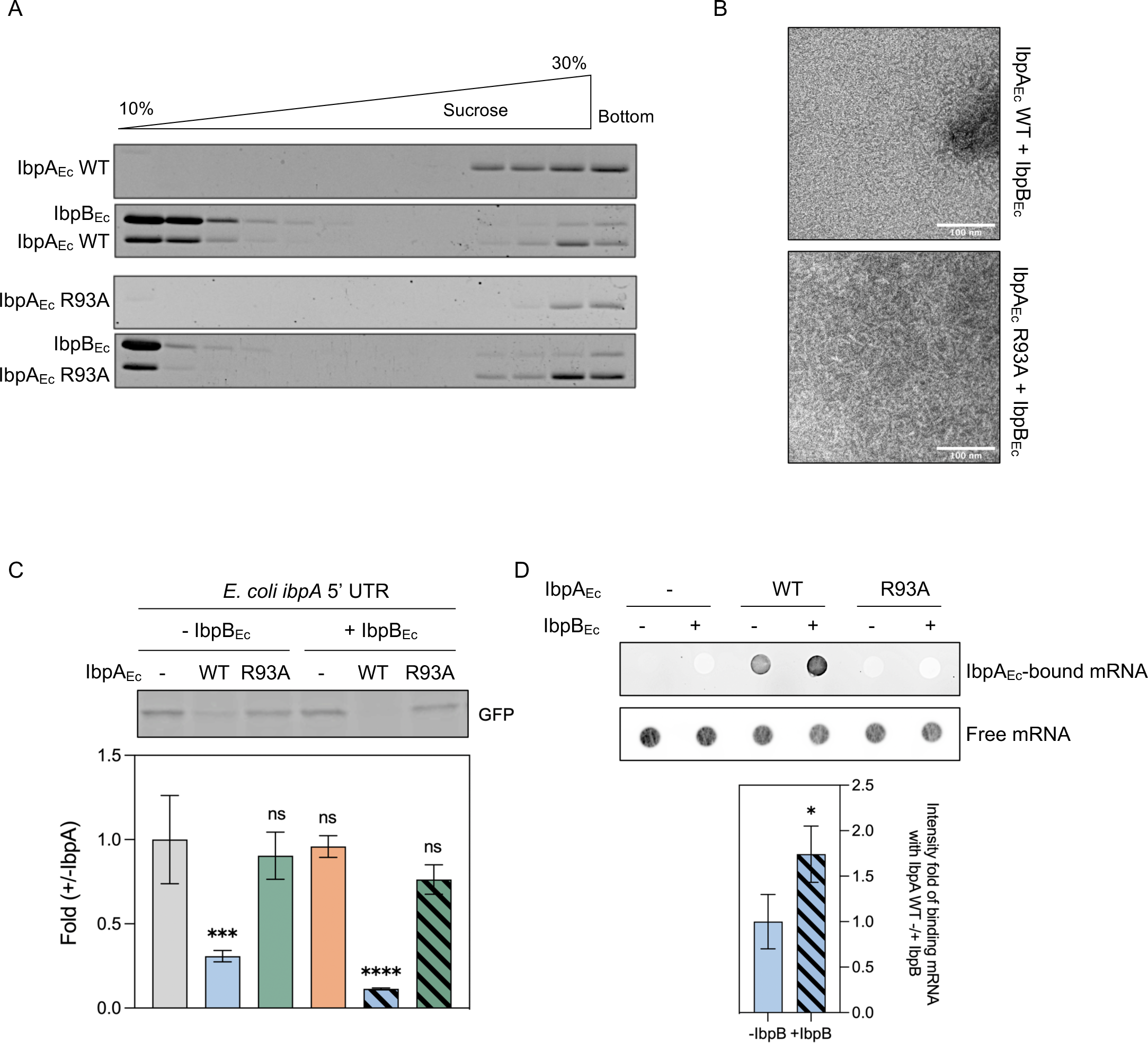
IbpB_Ec_ enhances the translation suppression ability of IbpA_Ec_-WT but exerts no effect on IbpA_Ec_-R93A. (*A*) Interaction between IbpA_Ec_ (WT or R93A) and IbpB_Ec_ assessed by SDG centrifugation. Purified IbpA_Ec_ in the presence or the absence of IbpB_Ec_ was subjected to 10%-30% (w/v) sucrose gradient solutions, as shown in Fig. 4. Collected fractions were analyzed by SDS-PAGE with 6 M urea to separate IbpA_Ec_ and IbpB_Ec_, and visualized by CBB staining. (*B*) Representative TEM images of purified IbpA_Ec_-WT (*upper*) and R93A (*lower*) in the presence of IbpB_Ec_. The scale bars represent 100 nm. (*C*) Effect of IbpA_Ec_ on the translation of the *ibpA_Ec_*5’ UTR-*gfp* reporter in the absence or presence of IbpB_Ec_. *Upper*: the fluorescence intensity of the translated GFP; *lower*: the fold-change in GFP translation level compared with that in the absence of both IbpA_Ec_ and IbpB_Ec_. (*D*) IbpA_Ec_-mRNA interaction in the presence or absence of IbpB_Ec_ assessed by filter binding assay. *Upper*: the biotin-labeled mRNA intensity detected by streptavidin Alexa Four 647 conjugate; *lower*: the quantified intensity fold of IbpA_Ec_-bound mRNA compared to that in the absence of IbpB_Ec_. All data represent the means (±S.D.) of three independent experiments and were analyzed by one-way ANOVA.

Furthermore, we assessed the effect of IbpB_Ec_ on IbpA-mediated translation suppression. We validated that IbpB_Ec_ had no impact on the translation of *ibpA* 5’ UTR-*gfp* in the PURE system, whereas it enhanced the IbpA-WT-mediated translation suppression (Fig. 6C). However, the absence of translation repression by IbpA_Ec_-R93A remained unchanged in the presence of IbpB_Ec_ (Fig. 6C). In the filter binding assay, we observed that the presence of IbpB_Ec_ enhanced the interaction of IbpA_Ec_-WT with the *ibpA* 5’ UTR mRNA, while IbpB_Ec_ did not alter the property of IbpA_Ec_-R93A to not bind to the mRNA (Fig. 6D).

## Discussion

After discovering the nonconventional function of IbpA_Ec_ in suppressing its own translation (18), there remain several fundamental questions to address. This study sheds light on some of the mechanisms underlying the unique function of IbpA that its paralog, IbpB, lacks. First of all, it is worth noting that not only IbpA_Ec_ but also other bacterial IbpAs, *C. neteri* and *V. harveyi* IbpAs (IbpA_Cn_ and IbpA_Vh_), have the function to suppress their own translation through the interaction with their 5’ UTR mRNAs, indicating that the IbpA-mediated self-regulation function is evolutionally conserved.

In exploring the difference between IbpA_Ec_ and IbpB_Ec_, we found that a cluster of positively-charged amino acids within the ACD, which is not conserved in IbpB_Ec_, is crucial for the translation suppression activity (Fig. 3B). Within the cluster, the R93, R97, and K98 residues in the IbpA_Ec_ are conserved in those in other bacterial IbpAs, suggesting a key role of these R/K residues in discriminating between IbpA and IbpB. Importantly, human sHsps (HspB4 and HspB5) also possess the conserved positively-charged residues, mutations of which are associated with diseases (20, 21). Although there is no evidence for the translation suppression activity in human sHsps, the positive-charge clusters in the ACD are likely of common importance for sHsps in both prokaryotes and eukaryotes.

Among the conserved positive-charge residues, the R93 position in IbpA_Ec_ has a unique role since it cannot be replaced by any of the other 19 amino acids, including Lys (Fig. S4). In contrast, R97 and R98 can be replaced by Lys, indicating that the cationic property of these residues is sufficient. It is noteworthy that this interconversion between K and R is largely the same trend observed in the positive-charge cluster in human sHsps (23). An AlphaFold2-predicted hexamer structure of IbpA_Ec_ revealed that R93, R97, and K98 are in close proximity to NTD (Fig. 3D), suggesting a potential interplay between the region and NTD. Unlike R97 and K98, which are located in a stable β7 sheet, R93 is situated in a flexible loop that bridges β6 and β7. Mutants of Y34 in a disordered loop of NTD, which is closest to R93 (Fig. 3D), abolish the translation suppression activity, unless the substitution is to aromatic residues (Fig. 3E) (see below). These results support the notion that the potential interaction between R93 and Y34 is responsible for the activity. Similar potential interactions are also apparent in the predicted IbpA_Cn_ and IbpA_Vh_ structures (Fig. S5), further supporting this notion. We note that IbpB_Ec_ also has an aromatic residue (F32) in the position corresponding to Y34 in IbpA_Ec_, which is consistent with the observation that the NTD-substituted chimera retained the self-suppression activity (Fig. S1A).

Why is it only possible for Arg to be in position 93, but not Lys? Although Arg and Lys are typically considered equivalent in their positive charge, there are notable differences in their properties. Among these differences, it is worth noting that Arg is abundant in RNA-binding proteins, as observed in previous studies (25–28). Furthermore, Arg is found more frequently at protein–protein interfaces than Lys (29, 30), which makes it a “stickier” amino acid. It is plausible that the “stickiness” of Arg, which could be involved in RNA binding, contributes to the ability of IbpAs to bind to mRNA, thus repressing self-translation.

When considering the irreplaceable nature of R93 and the importance of aromatic residues at the Y34 position in NTD, it is tempting to consider the possibility of a cation-π interaction in the mRNA binding activity. IbpA does not bind any mRNA indiscriminately, but rather seems to bind RNAs with secondary structure, such as RNATs (18). The cation-π interaction involving a specific RNA structure may be crucial to the nonconventional function of IbpA. Additionally, since the cation-π interaction is one of the driving forces for undergoing liquid-liquid phase separation (LLPS) (25–28), there is a possibility of LLPS in the IbpA-mediated translation suppression function, although our rationale for investigating the Tyr residue is based on an AlphaFold2-predicted structure that suggests intramolecular interaction. Nonetheless, a previous report on sHsps, which suggests HspB2 concentration-dependent LLPS formation in cells (31), implies a possible involvement of LLPS in the IbpA-mediated self-regulation activity.

We previously showed that IbpA_Ec_’s self-translation suppression relies on its interaction with the *ibpA* 5’ UTR mRNA using a gel shift assay (18). In this study, we developed a filter binding assay using purified protein and mRNA to analyze the IbpA_Ec_-mRNA interaction (Fig. 5 and 6). The filter binding assay revealed that IbpA_Ec_-R93A completely lost the ability to interact with the *ibpA* 5’ UTR mRNA, as expected from the in vivo reporter assay and the PURE system analysis. In contrast, the chaperone activity of IbpA_Ec_-R93A, as assessed by binding to heat-denatured proteins, showed only a small decrease. Furthermore, the chaperone activity of the R93 mutants in other bacteria (IbpA_Cn_ and IbpA_Vh_) was indistinguishable from that of the wild type. Therefore, the R93 mutation mainly abolishes the mRNA binding activity of IbpAs, indicating that the mRNA-binding activity of IbpAs and its chaperone activity to sequester denatured proteins are distinguishable.

The self-suppression function of IbpAs relies on the oligomer states. IbpA_Ec_-R93A tends to form higher-order oligomers as well as longer fibril-like structures (Fig. 4A, B). It is possible that the chaperone activity of IbpA_Ec_-R93A is slightly defective due to its propensity to form larger oligomers and fibril-like structures, which are known to be inactive as chaperones, as previously reported (17). The propensity of IbpA_Ec_-R93A to form larger-size oligomers and fibrils is consistent with its tendency to form oligomers in a broad size range, even after introducing the AEA mutation that inhibits IbpA_Ec_ oligomer formation. This indicates that the over-oligomerization of IbpA_Ec_-R93A is so robust that it can overcome the oligomerization-inability of the AEA mutation. In addition, even when IbpA_Ec_-R93A forms smaller oligomers in the presence of IbpB_Ec_, it is still repression-inactive (Fig. 6C, D). These findings provide evidence that the translation suppression function is independent of the oligomeric states.

Although the oligomerization behavior of IbpA_Ec_-R93A is different from that of wild-type, the oligomeric states of the other bacterial IbpA-R93 mutants were almost identical to those of the wild-type IbpAs (Fig. 4C, D). The AlphaFold2-based predicted structures indicate no apparent difference in dimer structures between the wild type and the R93 mutants for all IbpAs examined in this study (Fig. S8). This explains why the R93 mutants possess the same or similar chaperone activity. Collectively, these results strongly suggest that the defect of R93 mutants in the translation suppression activity through impaired mRNA binding is not caused by size variation in oligomers, implying a more complex mechanism behind the suppression activity.

The incorporation of IbpA-R93A into IbpA-WT abolished the WT-mRNA interaction (Fig. 5C-E). If we assume that the hetero-oligomer formation is not compromised in the R93A mutant, the incorporation of the mutants into the WT oligomer perturbs the WT-mediated mRNA binding. In contrast to R93A, IbpB_Ec_ can strengthen the interaction between IbpA_Ec_-WT and mRNA, possibly due to the reduced fibril formation of IbpA_Ec_-WT (Fig. 6B). Furthermore, the impaired suppression activity of IbpA_Ec_-R93A cannot be activated in the presence of IbpB_Ec_, even though R93A interacts with IbpB_Ec_ and forms more non-fibrous structures (Fig. 6A).

The AlphaFold2-predicted structures indicate that the monomeric structure of the R93A mutant is nearly identical to the wild type (Fig. S9), as expected for dimeric structures (Fig. S8). Previous analyses have already revealed that sHsp dimerization is promoted by the β6 loop of one ACD interacting with a β2 strand of the partner’s ACD, while oligomerization is induced by the interplay between the IXI/V motif in CTD and a β4/β8 groove of the neighboring dimer (4). The formation of hetero-dimers and hetero-oligomers of IbpA-IbpB has been found to be essentially the same as that of a single sHsp (16). All of these interacting regions are independent of the β6/β7-bridging loop region containing R93 (Fig. 3D), which ensures the interplays of IbpA-R93A with IbpA-WT and IbpB.

Overall, this study demonstrates that IbpA-mediated self-suppression function is conserved in bacteria, and that the cationic residues in ACD, particularly R93, are essential for this function. Since the Arg93 mutations in this region have little effect on chaperone activity, the conservation of Arg93 in bacterial IbpAs implies that the translation suppression function would be critical for the cellular function of IbpAs, in addition to the sequestrase function as a chaperone.

## Materials and Methods

### Plasmid construction and cloning

Constructions of pBAD30-*ibpA_Ec_* 5’ UTR-*gfp* and pCA24N-*ibpA_Ec_* were reported previously (18). The *ibpAB_Ec_* chimeras were produced by standard cloning procedures and Gibson assembly. Single mutations in pCA24N-*ibpA_Ec_* were achieved by site-directed mutagenesis. The pCA24N-*ibpA_Cn_* and pCA24N-*ibpA_Vh_*were individually constructed by sub-cloning of *ibpA_Cn_* and *ibpA_Vh_* DNA fragments into pCA24N using Gibson assembly. The *ibpA_Cn_*and *ibpA_Vh_* DNA fragments were amplified from pET3a-*ibpA_Cn_*and pET3a-*ibpA_Vh_* plasmids, which were kindly provided by Dr. Krzysztof Liberek (15). The IbpA_Cn_-R93A and IbpA_Vh_-R94A mutants were also constructed individually by site-direct mutagenesis from pCA24N-*ibpA*_Cn_ and pCA24N-*ibpA*_Vh_. The pBAD30-*ibpA_Cn_* 5’ UTR-*gfp* and the pBAD30-*ibpA_Vh_*5’ UTR-*gfp* were generated by sub-cloning of the *ibpA_Cn_* 5’ UTR and the *ibpA_Vh_* 5’ UTR DNA fragments into pBAD30-*gfp*, respectively. The DNA fragments of the *ibpA_Cn_* and the *ibpA_Vh_* 5’ UTRs were amplified from DNA oligos commercially produced by Integrated DNA Technologies (USA). The sequence information of the *ibpA_Cn_* and the *ibpA_Vh_* 5’ UTRs was obtained from National Library of Medicine NBRC 105707 and NBRC 15634. All plasmids were amplified in *E. coli* DH5α. Primers used for cloning and mutagenesis were shown in Table S2.

### Reporter Assay

The pBAD30 plasmids, as reporter plasmids, were transformed or co-transformed with pCA24N plasmids into *E. coli* BW25113 wild-type and Δ*ibpAB,* where *E. coli* chromosomal *ibpAB* operon was deleted as reported previously (18). After pre-culture in LB medium at 37°C with 180 rpm overnight, the cells were diluted to fresh LB medium with the induction of 2×10^-4^ % arabinose until growing to OD_660_ of 0.2 ∼ 0.3. The cells containing only pBAD30 plasmids were incubated for more than 30 min, while the cells possessing both pBAD30 and pCA24N plasmids were further induced with 0.1 mM isopropyl-β-D-thiogalactopyranoside (IPTG) for 30 min to achieve the overexpression of exogenous IbpAs. Next, the cells were harvested and treated with the same-volume of 10% TCA to precipitate proteins. After incubation on ice for 15 min and centrifugation at 20,000 g for 3 min at 4°C, the pellets were washed by ice-cold acetone and then centrifugated again. After 2 times of acetone washing, 1X SDS sample buffer (0.25 M Tris-HCl pH 6.8, 5% SDS, 5% (w/v) sucrose, 0.005% (w/v) bromophenol blue, and 5% (w/v) 2-mercaptoethanol) was applied to dissolve the pellets, and then the samples were incubated at 37°C for 15 min. The SDS-treated samples were applied to 12% polyacrylamide SDS gels and then transferred from the gels to PVDF membranes based on standard immunoblotting procedures. The membranes were blocked with 1% skim milk in TBS-T buffer (20 mM Tris-HCl pH 7.5, 137 mM NaCl, and 0.2% (w/v) Tween20). Mouse anti-sera against GFP (mFx75, Wako) or rabbit anti-sera against FtsZ (kindly provided by Dr. Shinya Sugimoto from Jikei Medical University) was used as the primary antibody with the dilution of 1:10,000. The secondary antibody was HRP-conjugated anti-mouse or -rabbit IgG (Sigma-Aldrich) used in the same dilution factor. Subsequently, the samples were visualized by Dual Chemiluminescent substrates (Millipore_®_) and detected by a LAS 4000 mini imager (Fujifilm).

### Protein purification

To purify IbpA_Ec_ and its mutants (R93A, AEA, and R93A/AEA), *E. coli* BW25113ΔAB cells at OD_660_ of 0.5 were used to overexpress IbpA_Ec_-WT, -R93A, - AEA and -R93A/AEA with the induction of 0.5 mM IPTG at 37°C for 4 h. The harvested cells were lysed by sonication (Branson) in buffer A (50 mM Hepes-KOH pH 7.6, 10% glycerol, 1 mM dithiothreitol, and 0.1 M KCl). The following anion exchange chromatography was performed as described previously (18). The purification of soluble IbpA_Ec_-R93A/AEA required one more Q-Sepharose (Cytiva) chromatography step with elution by a pH gradient from buffer A to buffer B (50 mM critic acid-NaOH pH 5.0, 10% glycerol, 1 mM DTT and 0.1 M KCl). His-IbpB_Ec_ was purified with Ni-NTA agarose upon the denaturation by 6 M urea. Purification of IbpA_Cn_ WT and R93A expressed in *E. coli* was the same as that of IbpA_Ec_. To purify IbpA_Vh_ WT and R94A, pET3a-*ibpA_Vh_* or *ibpA_Vh_*R94A was overexpressed in *E. coli* BL21(DE3) cells at OD_660_ of 0.5 with the induction of 10 µM IPTG at 28°C for 20 hours. The harvested cells were lysed by sonication in buffer C (50 mM Tris-HCl pH 7.5, 10% glycerol, and 1 mM DTT). IbpA_Vh_ was soluble in *E. coli* cells, so supernatants of the lysates were collected by low-speed centrifugation (10,000 g, 10 min, 4°C) to remove *E. coli* inclusion bodies including endogenous IbpA_Ec_. Then, one more centrifugation with high speed (30,000 rpm, 30 min, 4°C) was carried out to remove the pellet of membrane vesicles and ribosomal particles. Next, the supernatants were applied to QAE resin (Toyopearl®, Tosoh), and the flowthrough fractions were collected since most of the native IbpA_Vh_-WT and -R94A were found in these fractions. Then the flowthroughs were applied onto fresh QAE resin after being dialyzed against buffer D (50 mM Tris-HCl pH 7.5, 10% glycerol, 1 mM DTT, and 6 M urea). The denatured IbpA_Vh_ was eluted by increasing salt concentration to 200 mM NaCl in buffer D. The IbpA_Vh_-containing fractions were then dialyzed against buffer D and applied to another round of QAE chromatography with the elution of decreasing pH from buffer D to buffer E (50 mM critic acid-NaOH pH 5, 10% glycerol, 1 mM DTT and 6 M urea). Finally, urea was gradually removed by dialysis against buffer F (50 mM Tris-HCl pH 7.5, 10% glycerol, 1 mM DTT and 0.1 M KCl) from buffer F containing 4 M urea, 2 M urea, 1 M urea to buffer F without urea. Purification of firefly luciferase was performed as described previously (32). Protein concentrations were determined by the Bradford method with standard BSA. All concentrations of IbpA in this study were in dimeric units.

### Reconstituted cell-free translation

PUREfrex^®^ (GeneFrontier) with Cy5-labeled tRNA^fMet^ was used to express RNA templates produced by CUGA^®^7 in vitro transcription kit (Nippon Gene) in the presence or the absence of purified IbpAs or the mutants (1 µM). The reaction mixtures were incubated at 37°C for 2 h, then mixed with the same volume of 2x SDS sample buffer (0.5 M Tris-HCl pH 6.8, 10% SDS, 10% (w/v) sucrose, 0.01% (w/v) bromophenol blue and 10% (w/v) 2-mercaptoethanol), and boiled at 95°C for 5 min. The mixtures were separated by SDS-PAGE, and visualized by an Amersham™ Typhoon™ scanner (Cytiva), and finally quantified by Multi gauge software (Fujifilm).

### Sucrose density gradient centrifugation

To investigate the oligomeric states of the purified IbpAs and the mutants, the proteins (3 µM) in buffer G (50 mM Hepes-KOH pH 7.6, 0.1 M KCl, 5 mM DTT and 20 mM Mg-acetate) were applied onto a 11 ml 10-30% (w/v) sucrose gradient in buffer G and then ultracentrifuged with a Beckman SW41Ti rotor (35,000 rpm, 4°C, 80 min). The centrifuged samples were collected from the top to the bottom by a fractionator (BioComp). Then, the top 6 fractions, the bottom 6 fractions and the aggregates attached to the tube bottom were analyzed by SDS-PAGE, and detected by standard western blotting procedures using rabbit anti-sera against IbpA (Eurofin) as primary antibody and HRP-conjugated anti-rabbit IgG as secondary antibody (Sigma-Aldrich). For IbpA_Cn_ and IbpA_Vh_ detection, Coomassie Brilliant Blue (CBB) was used for visualization.

To investigate the interaction of IbpAs with substrate proteins, the purified IbpAs (12 µM) and the purified luciferase (3 µM) in buffer G were mixed and incubated at 50°C for 30 min. After that, the mixtures were applied onto a 10-50% (w/v) sucrose gradient and then centrifuged as described above. The protein distributions were verified by SDS-PAGE and visualized by CBB staining.

To investigate interaction with IbpB_Ec_, the mixtures of IbpA_Ec_ (3 µM) and IbpB_Ec_ (7 µM) in buffer G were incubated at room temperature for 30 min and then applied onto a 10-30% (w/v) sucrose gradient. Centrifugation and fraction collection were performed as above. The fractions were analyzed by SDS-PAGE in the presence of 6 M urea followed by CBB staining.

### Transmission electron microscopy

Purified IbpA_Ec_-WT or -R93A (2 µM) in the absence or presence of IbpB_Ec_ (4.8 µM) was applied on carbon-coated copper grids. The samples were allowed to absorb for 1 min before negatively stained with 1% methylamine tungstate at pH 7 for 1 min. The staining was repeated twice. The observation was performed with a JEOL 1400 Plus electron microscopy.

### Filter binding assay

The *ibpA* 5’ UTR*-gfp* mRNA produced from CUGA^®^7 in vitro transcription kit (Nippon Gene) was attached to a 3’-terminal biotinylated nucleotide using Pierce™ RNA 3’ End Biotinylation kit (Thermo Scientific). The biotin-labeled mRNA (0.1 µM) was incubated with different ratio of IbpA in buffer H (100 mM sodium phosphate buffer pH 7.5, 0.1 M NaCl, 5 mM EDTA, 5 mM DTT and 10% glycerol) at room temperature for 30 min, after which the mixtures were fixed with 1% formaldehyde for 10 min followed by an addition of 0.25 M glycine to stop the cross-linking reactions in 5 min. A nitrocellulose membrane (Amersham™ Protran™ 0.2 µM NC, GE Healthcare, Life Sciences) was pre-soaked in buffer H and then overlaid on a positively-charged nylon membrane (BrightStar™-Plus, Invitrogen). The protein-mRNA mixtures were applied onto a 96-well slot-blot apparatus and then filtered through the double membranes by vacuum. The protein-mRNA complexes were trapped in the top nitrocellulose membrane, while the free mRNA samples passed through the nitrocellulose membrane and were caught by the bottom nylon membrane. Finally, biotin-labeled mRNAs were detected with streptavidin-HRP (Thermo Scientific) according to the protocol prepared for Chemiluminescent Nucleic Acid Detection Module kit (Thermo Scientific). To examine the interaction of IbpA_Ec_-WT with the mRNA in the presence of IbpA_Ec_-R93A, the mixture containing the mRNA (0.1 µM), IbpA_Ec_-WT (1 µM) and different ratios of IbpA_Ec_-R93A was applied to the filter assay, and the biotin-labeled mRNA was detected by streptavidin Alexa Fluor™ 647 conjugate (Invitrogen™). To investigate the effect of IbpB_Ec_ on IbpA_Ec_-mRNA interaction, the mixture containing the mRNA (0.1 µM), IbpA_Ec_ (1 µM) and IbpB_Ec_ (2.4 µM) was applied to the filter assay, and the biotin-labeled mRNA was detected by streptavidin Alexa Fluor™ 647 conjugate (Invitrogen™).

### Statistical Analysis

One-way ANOVA was used for calculating statistical significance. All experiments were conducted at least three times independently, and the mean values ± standard deviation (SD) were represented in the figures.

### Data availability

Data in this manuscript have been uploaded to the Mendeley Dataset public repository (https://doi.org/10.17632/gj92kb2wdd.1).

## Acknowledgements

We thank Krzysztof Liberek for the plasmids harboring *ibpA_Cn_* and *ibpA_Vh_*, Shinya Sugimoto for the anti-FtsZ antibody, Keiko Ikeda for electron microscopy, Naohiko Shimada and Atsushi Maruyama for CD measurement, the Biomaterials Analysis Division, Open Facility Center at Tokyo Tech for DNA sequencing. This work was supported by MEXT Grants-in-Aid for Scientific Research (Grant Numbers JP26116002, JP18H03984, and JP20H05925 to HT, JP22K14860 to TM) and JST SPRING Grant number JPMJSP2106 to YC.

## Author Contributions

Y.C. performed experiments; Y.C., T.M., and H.T. conceived the study, designed experiments, and analyzed the results; H.T. supervised the entire project; Y.C. T.M. and H.T. wrote the manuscript.

