## Supplementary Fig S1-S9, Supplementary Table S1, S2 for "Self-regulatory function of bacterial small heat shock protein IbpA through mRNA binding is conferred by a conserved arginine"

**This PDF file includes:**

Figures S1 to S9
Table S1, S2
SI References

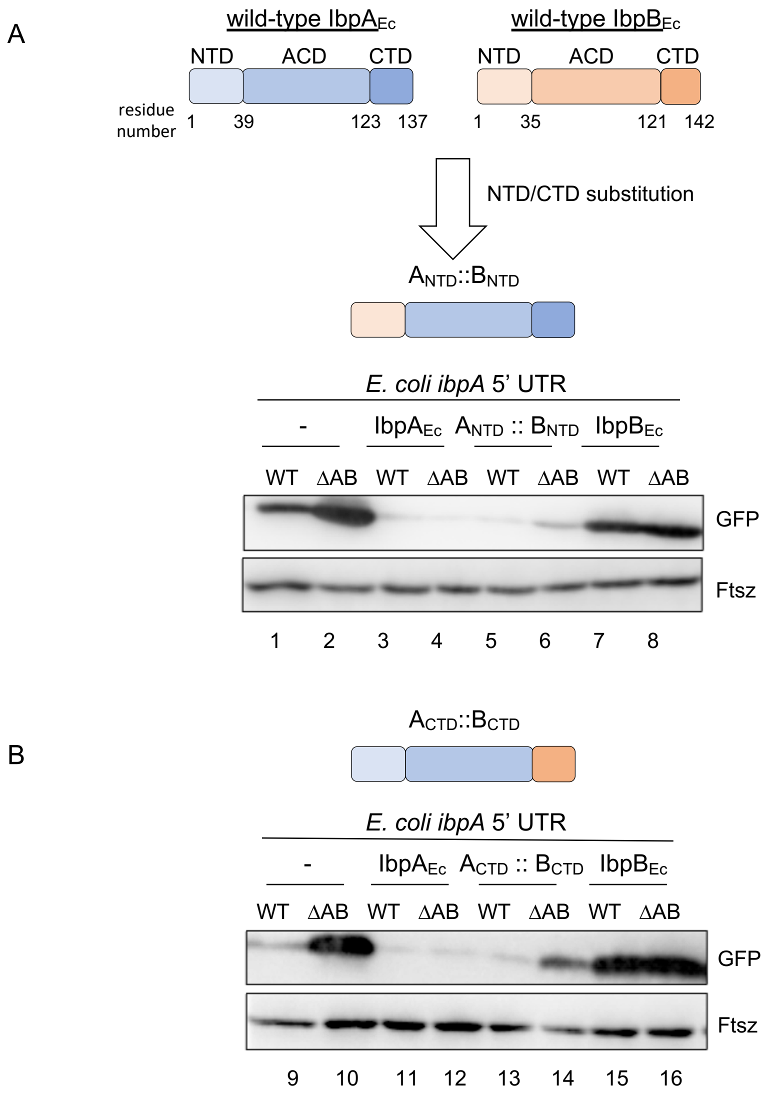


**Fig. S1. Effects of IbpA$_{Ec}$/IbpB$_{Ec}$ chimeras in the N-terminal domain (NTD) or C-terminal domain (CTD) on the IbpA$_{Ec}$-mediated translation suppression.**
*Upper*: a schematic representation of IbpA$_{Ec}$/IbpB$_{Ec}$ chimeras in NTD (*A*) or CTD (*B*), where IbpA$_{Ec}$ and IbpB$_{Ec}$ are denoted by blue and orange, respectively. *Lower*: Western blotting analysis to evaluate the effects of the NTD (*A*) and CTD (*B*) chimeras on the level of the GFP reporter translation in *E. coli* BW25113 strains (WT: BW25113 wild-type strain; ΔAB: *ibpAB* operon-deleted BW25114 strain). The constitutive expression level of FtsZ is shown as a control. Anti-GFP and anti-FtsZ antibodies were used for the detection.

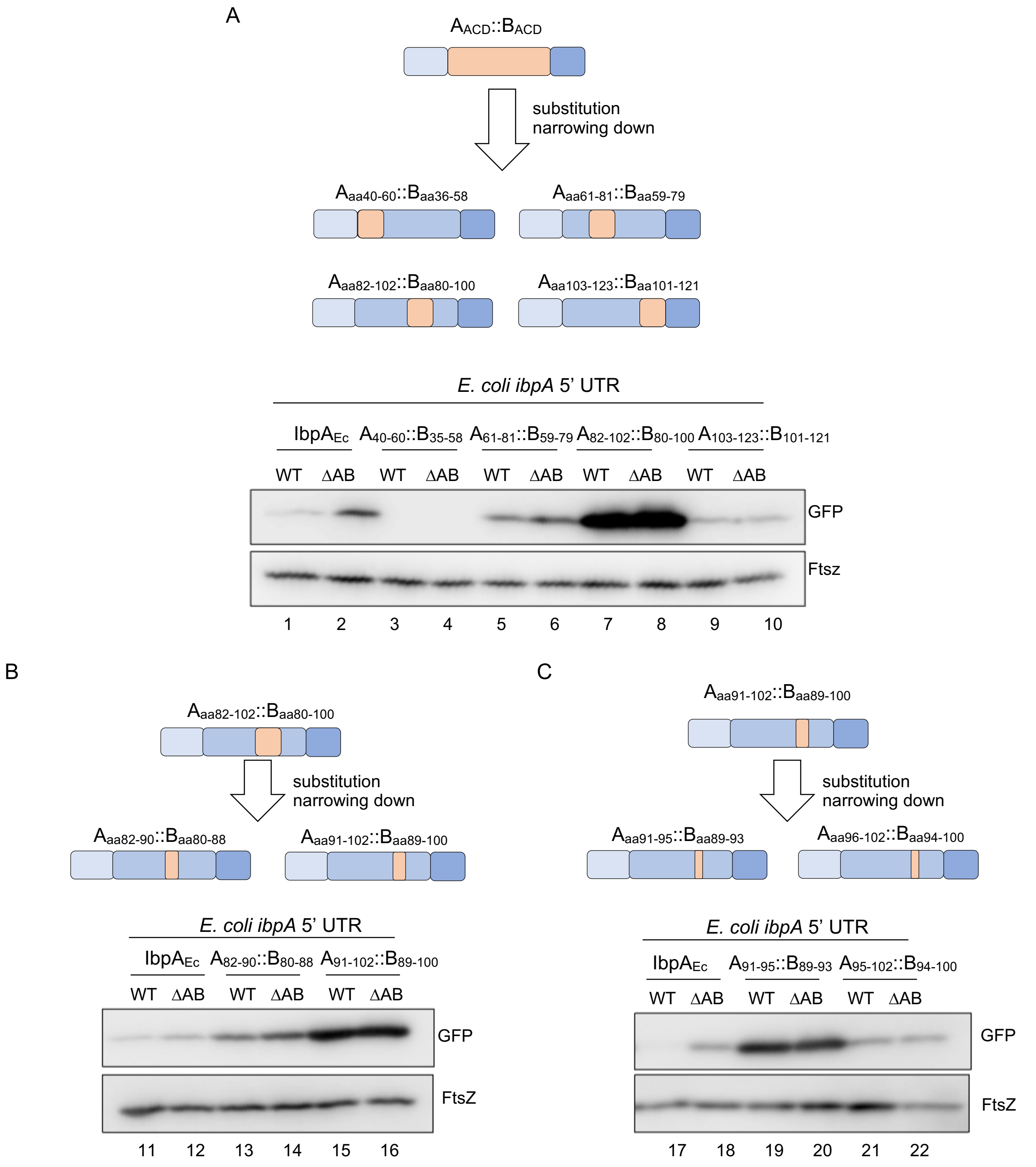


**Fig. S2. Systematic mutagenesis introduced in the IbpA$_{Ec}$-ACD to identify a region crucial for IbpA$_{Ec}$-mediated translation suppression.**

Following Fig. S1, IbpA$_{Ec}$/IbpB$_{Ec}$ chimeras further subdivided within the ACD are analyzed. The reporter assay method, including western blotting, is the same as that used in Fig. 1C, D and Fig. S1. (*A*) The ACD of IbpA$_{Ec}$ was subdivided into four parts, and the corresponding portions of IbpB$_{Ec}$ were individually substituted for each part. The A$_{82\text{-}102}$::B$_{80\text{-}100}$ chimera (IbpA$_{Ec}$ residues from 82 to 102 were replaced with IbpB$_{Ec}$ residues from 80 to 100) was found to be suppression-inactive compared to the other three chimeras and the wild-type IbpA$_{Ec}$. (*B*, *C*) The substitution regions of the suppression-defect chimera A$_{82\text{-}102}$::B$_{80\text{-}100}$ were further narrowed down until the IbpA$_{Ec}$ 91-95 region was identified as being critical for translation suppression.

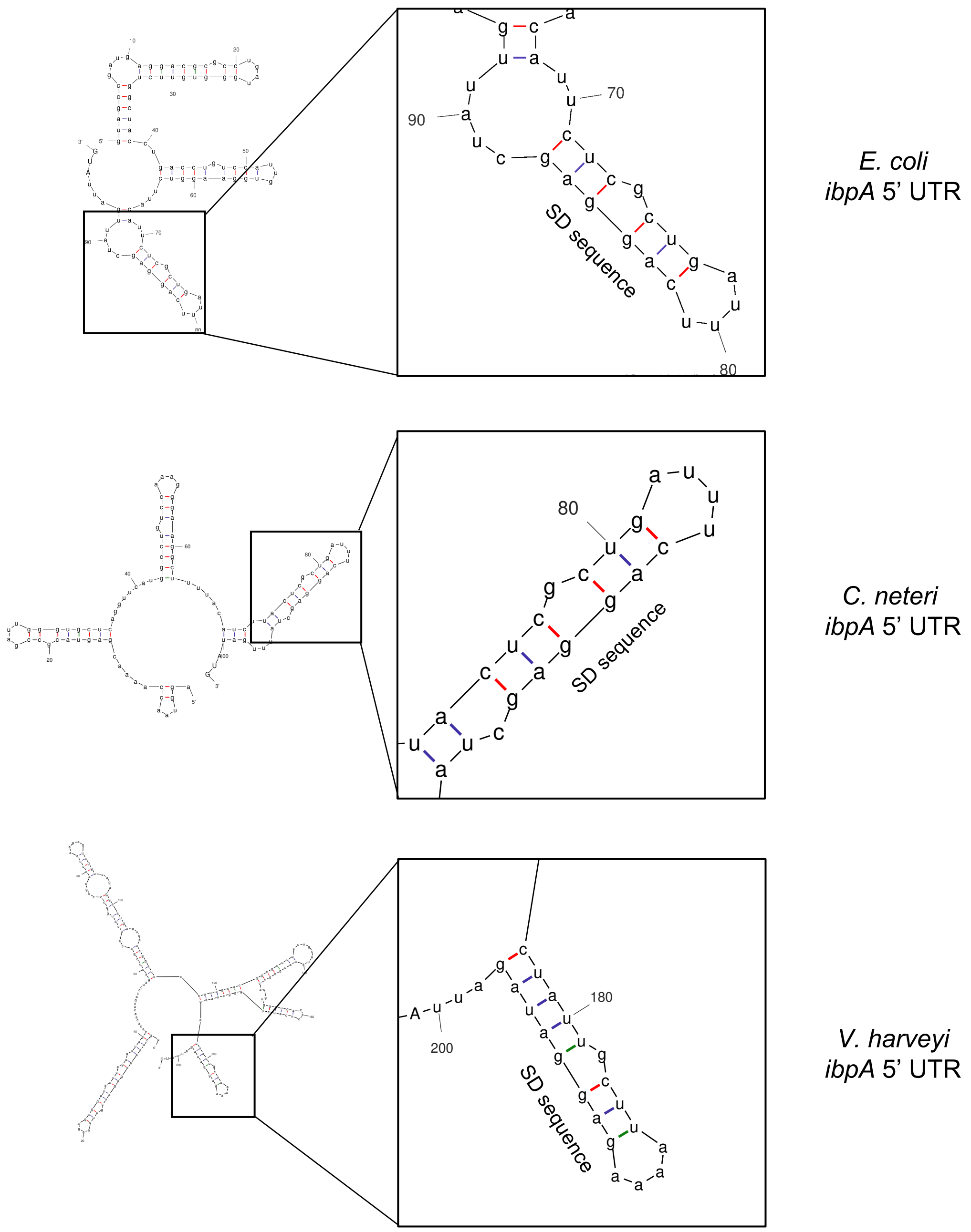


**Fig. S3. Prediction of RNAT-like structures in the 5' UTR of bacterial *ibpA* mRNAs.**

The secondary structures of the 5' UTR mRNAs of *ibpA*$_{Ec}$, *ibpA*$_{Cn}$, *and ibpA*$_{Vh}$, whose sequences are listed in Table S1, were predicted using UNAFold (http://www.unafold.org/). The Shine-Dalgarno sequences are highlighted.

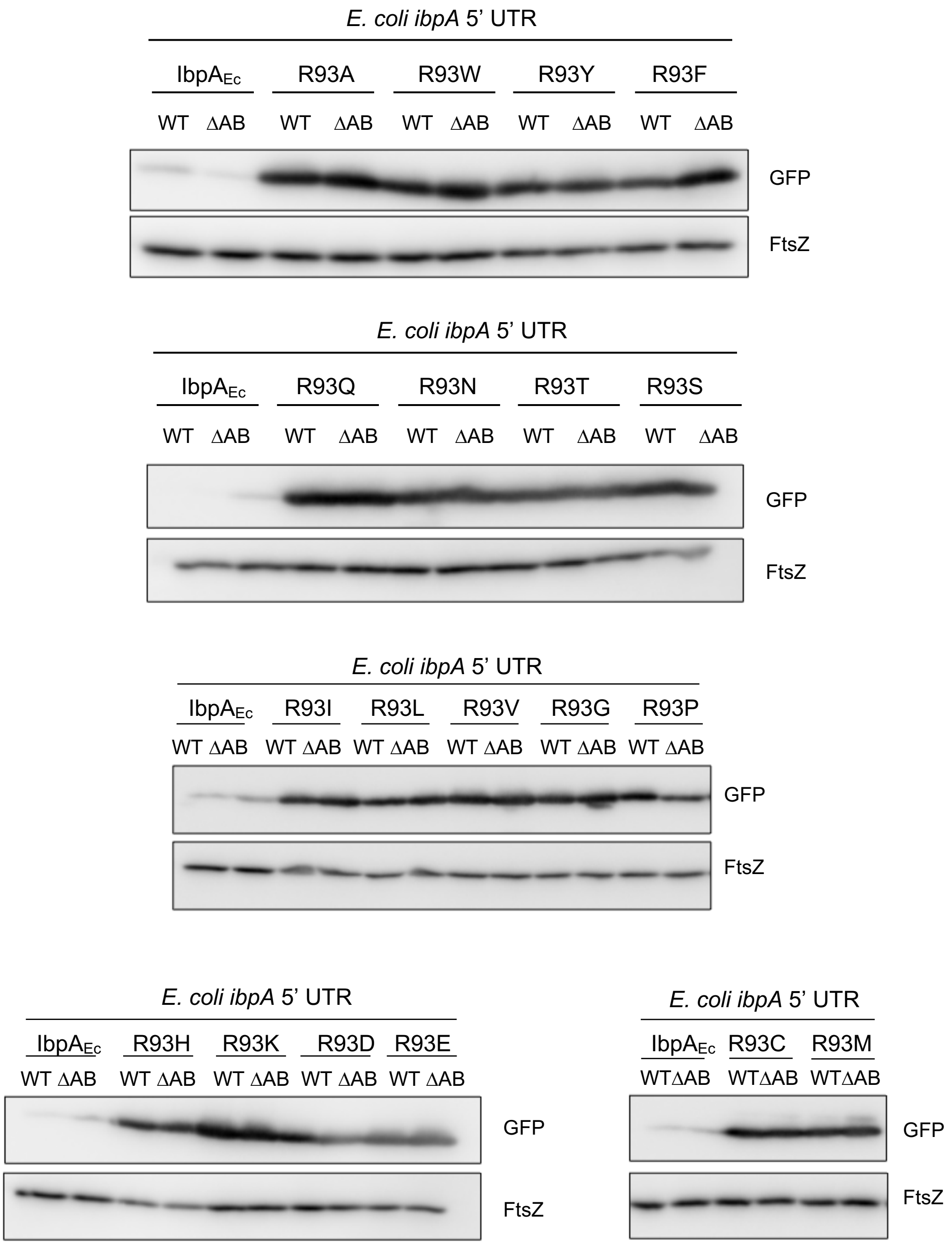


**Fig. S4. Mutations of the IbpA$_{Ec}$-R93 residue to other 19 amino acids.**

The arginine 93 in IbpA$_{Ec}$ was individually mutated to 19 other amino acids. The reporter assay method, including western blotting, is the same as that used in Fig. 1C, D and Fig. S1 and S2.

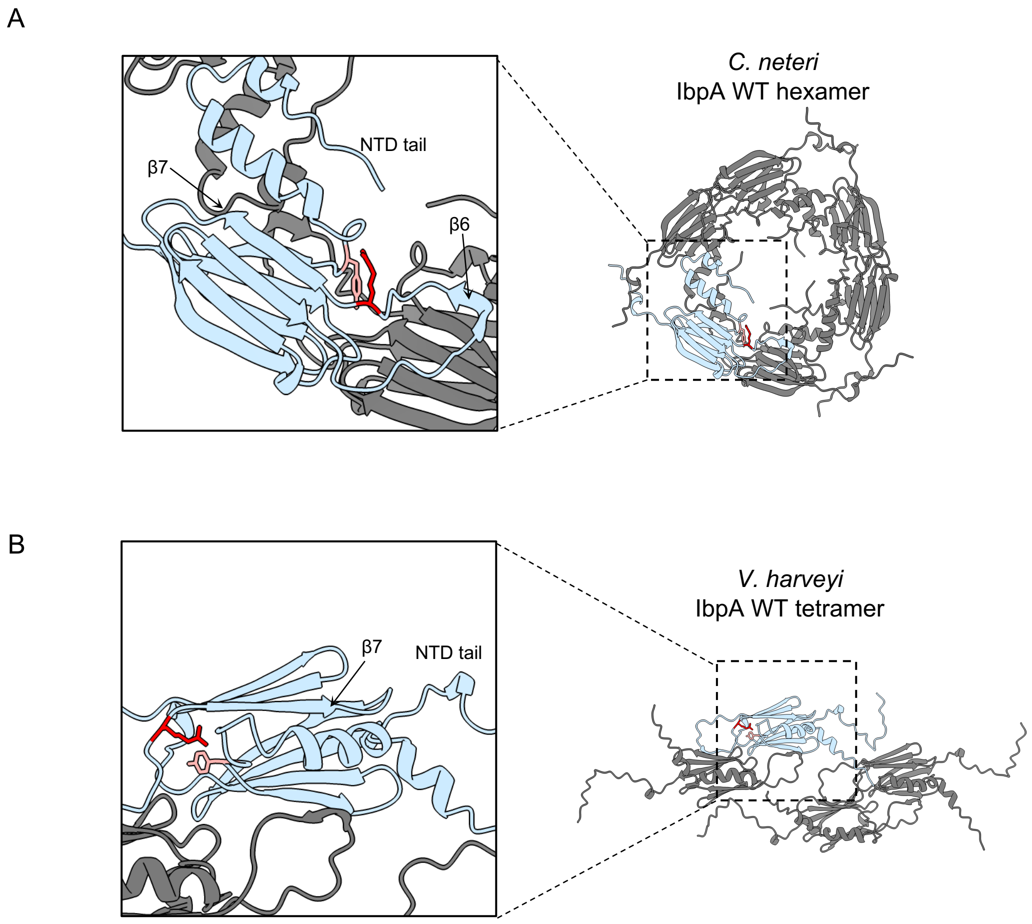


**Fig. S5. Potential interaction between conserved Y34 and R93 based on AlphaFold2-predicted IbpA structures.**

AlphaFold2-predicted structures of $\mathrm{IbpA_{Cn}}$-WT hexamer (*A*) and $\mathrm{IbpA_{Vh}}$-WT tetramer (*B*) using MMseqs2 (1), are shown. One of the subunits is colored light blue for clarity. Enlarged images indicate that Y34 and the Arg residues corresponding to $\mathrm{IbpA_{Ec}}$-R93 ($\mathrm{IbpA_{Cn}}$-R93 and $\mathrm{IbpA_{Vh}}$-R94) are colored pink and red, respectively.

A

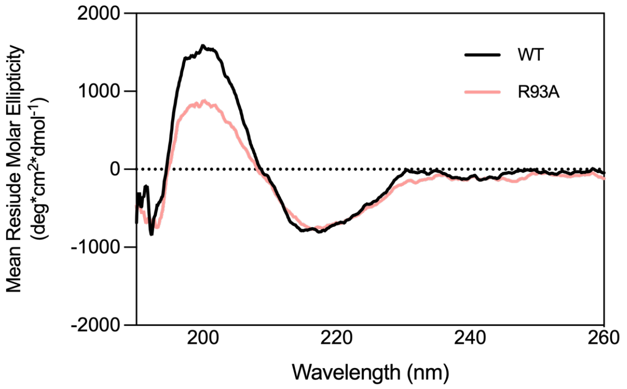


B

| % | α-Helix | Anti-parallel | Parallel | β-Turn | random coil |
|---|---|---|---|---|---|
| WT | 0 | 42 | 0 | 13.3 | 44.7 |
| R93A | 0 | 41.9 | 0 | 13.2 | 44.9 |

**Fig. S6. Circular Dichroism (CD) spectra of purified IbpA proteins.**
(*A*) The far-UV CD spectra of $IbpA_{Ec}$-WT, R93A (1.0 mg/ml) were measured using a JASCO J-820 spectropolarimeter (Japan) at 20 °C in a buffer containing 50 mM sodium phosphate pH 7.6. The measurement was performed using a 1 mm path-length cell, and each spectrum was an average of 10 accumulations. (*B*) The estimation of the secondary structure contents of IbpAs was calculated using BeStSel https://bestsel.elte.hu/index.php (2).

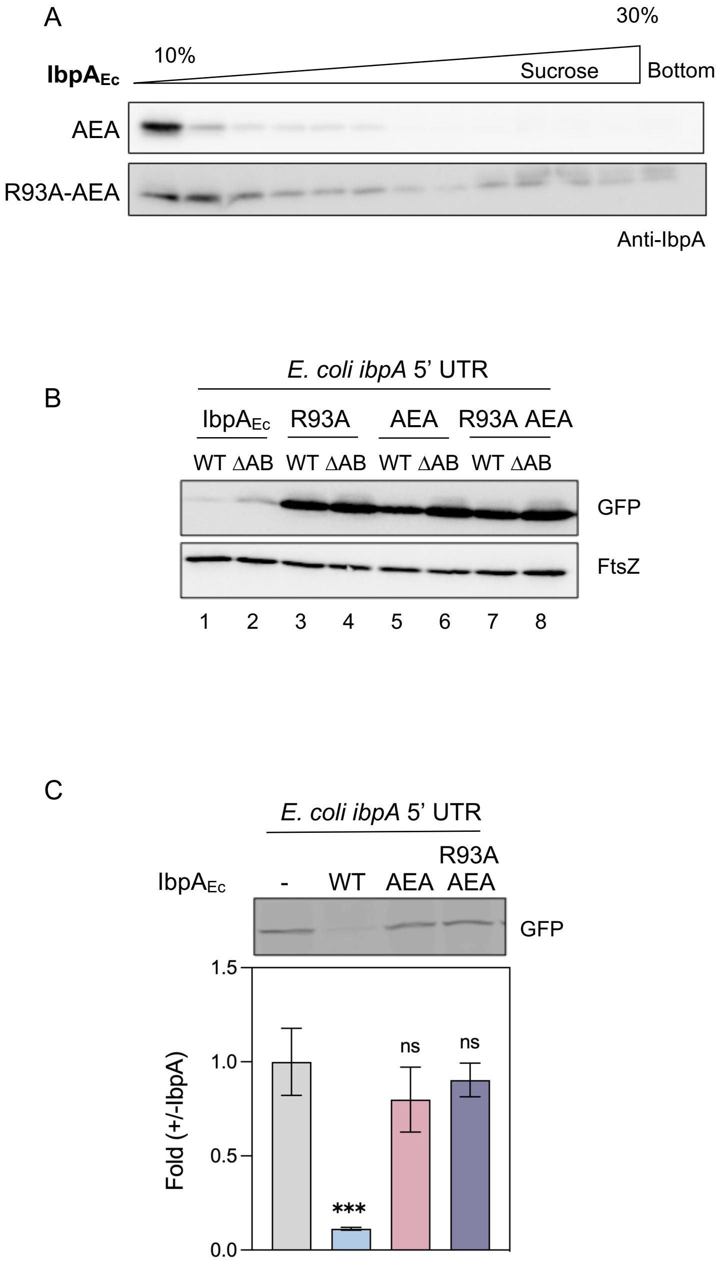


**Fig. S7. Characterization of the IbpA$_{Ec}$-R93A/AEA mutant.**

(*A*) The oligomeric states of IbpA$_{Ec}$-AEA and R93A/AEA mutants were evaluated using SDG centrifugation, as described in Fig. 4. (*B*) The effects of IbpA$_{Ec}$-AEA and R93A-AEA on the translation level of the *ibpA$_{Ec}$* 5' UTR-*gfp* in vivo were analyzed using a reporter assay method as shown in Fig. 1C, D and Fig. S1. (*C*) The effects of IbpA$_{Ec}$-AEA and R93A/AEA on the translation of the *ibpA$_{Ec}$* 5' UTR-*gfp* reporter in the PURE system were investigated using the method shown in Fig. 1E.

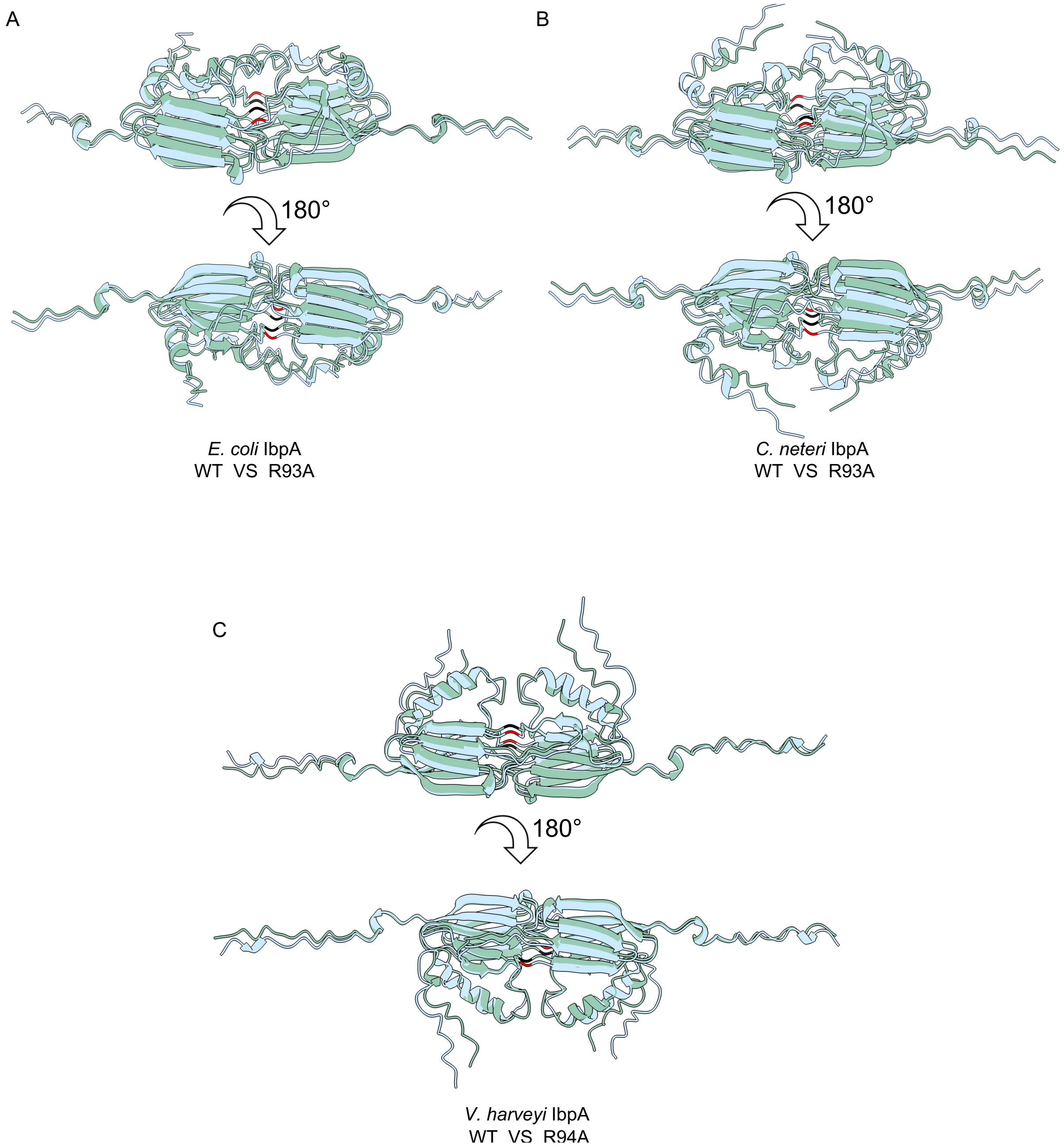


**Fig. S8. Predicted dimer structures of IbpA-WT and the R93 mutants.**
Predicted dimer structures of IbpA WT (*blue*) and the R93 mutants (*green*) in *E. coli* (*A*), *C. neteri* (*B*), and *V. harveyi* (*C*). The prediction was carried out using AlphaFold2 using MMseqs2 (1). The Arg93 (and the equivalent Arg residues) in the WT and Ala93 (and the equivalent Ala residues) in the R93A mutants are colored red and black, respectively.

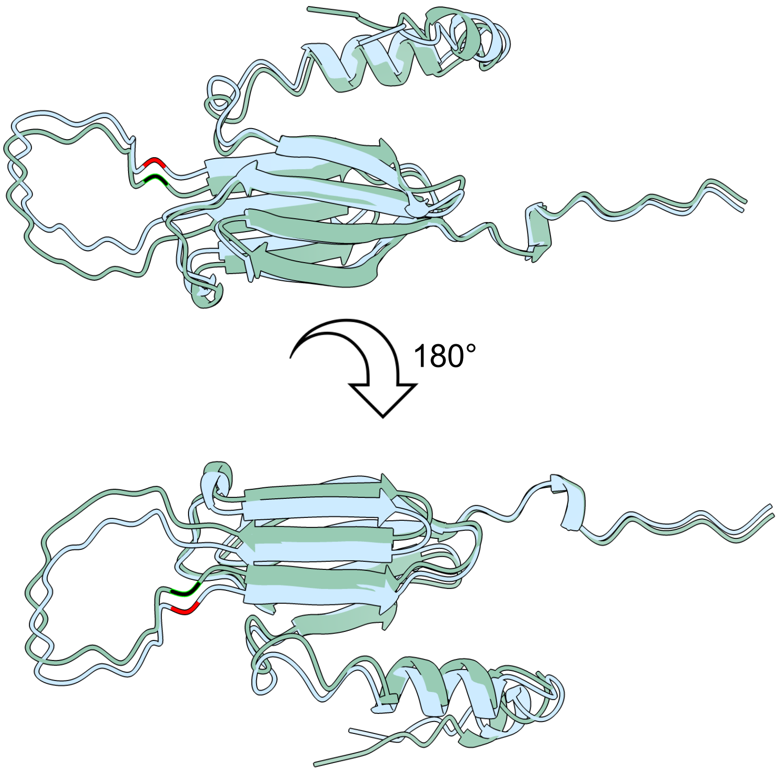


**Fig. S9. Predicted monomer structures of IbpA$_{Ec}$-WT and R93A.**

Predicted monomer structures of IbpA$_{Ec}$-WT (*blue*) and R93A (*green*) were obtained using AlphaFold2 with MMseqs2 (1). The R93 in WT and A93 in the R93A mutant are colored red and black, respectively.

Table S1. 5’ UTR sequences of IbpAs examined in this study.

| Name | Sequence |
|---|---|
| *$ibpA_{Ec}$ 5’ UTR* | gtagccgatgaggacgcgcctgatgggtgttctggctacctgacctgtccattg<br>tggaaggtcttacattctcgctgatttcaggagctattgattATG |
| *$ibpA_{Cn}$* 5’ UTR | aggtaaccaaaacgagtacgccgattgggtgctcaggttcatggcctgtccaaa<br>gggaaggcttttacatcttactcgctgatttcaggagctatttgatATG |
| *$ibpA_{Vh}$* 5’ UTR | cgctaaacgagatcgctcaagagaggatactcagtattagcacacaacttgaccc<br>tagattcgacatcatgtccccgcacctaaagagtaggtcagaggaatgaatcatcgg<br>atcccttagctcggcgttttcttgcataaatgcgccacgtggatgaccctcatctgggcta<br>agactattgcttaaaagaggatagattATG |

Table S2. Primers used in this study.

| Primer name | Sequence |
|---|---|
| pCA_ibpB_Fw | GAGGAGAAATTAACTATGCGTAACTTCGATTTATCCCCACTG |
| pCA_ibpB_Rv | GCTGCAGGTCGACCCTTAGCTATTTAACGCGGGACGTTCGC |
| ibpA-N::ibpB-N_Rv | GGTTTTCGTCTACCAGTTCgtacggcgggaagctctggctttc |
| ibpB-N_ibpA_alpha_Fw | gagcttcccgccgtacGAACTGGTAGACGAAAACCATTACCGC |
| ibpA_alpha_Rv | GCGTTCGAGATCGATATACAGCAAACCATTTACC |
| ibpA_alpha_ibpB_Cd_Fw | TGTATATCGATCTCGAACGCAATGAGCCTGAACCCATCGCAG |
| ibpB-C::ibpA-C_Fw | CATATTGATTTAATTCGTGTGATTCCGGAAGCGAAAAAACCG |
| ibpB-C::ibpA-C_Rv | CGCTTCCGGAATCACACGAATTAAATCAATATGCAGTAAACCG |
| ibpB-N::ibpA-N_Rv | cgtcgcttttctcaatgttAACGTTATACGGAGGGTAGCCGCC |
| ibpA-N_ibpB_alpha_Fw | CCCTCCGTATAACGTTaacattgagaaaagcgacgataacc |
| ibpB-C::ibpA-C_Fw | CATATTGATTTAATTCGTGTGATTCCGGAAGCGAAAAAACCG |
| ibpB-C::ibpA-C_Rv | CGCTTCCGGAATCACACGAATTAAATCAATATGCAGTAAACCG |
| pCA_ibpA_alpha1/21::B_ins_Fw | CGGCTACCCTCCGTATAACGTTgagaaaagcgacgataaccactaccgc |
| pCA_ibpA_alpha1/21::B_ins_Rv | CTGGGCGGTAATTTCCAGatcttcctgacggaaacctgccagcg |
| pCA_ibpA_alpha1/21::B_vec_Fw | GGAAATTACCGCCCAGGATAATCTGCTGGTGGTGAAAGGTGCTCACGC |
| pCA_ibpA_alpha1/21::B_vec_Rv | TACGGAGGGTAGCCGCCATTACTCTGGCTCTGGTTGTTTTCTAAGTG |
| pCA_ibpA_alpha22/42::B_ins_Fw | GTGGCTGGTTTTGCTGAGAGCGAAttagagattcaactggaaggtacg |
| pCA_ibpA_alpha22/42::B_ins_Rv | GGTACAGATAGGTGCGCTCttcttttggctgctccggcgtg |
| pCA_ibpA_alpha22/42::B_vec_Fw | GCGCACCTATCTGTACCAGGGCATCGCTGAACGCAACTTTGAAC |
| pCA_ibpA_alpha22/42::B_vec_Rv | CAGCAAAACCAGCCACAGCGATAGCAATGCGGTAATGGTTTTCGTC |
| pCA_ibpA_alpha43/63::B_ins_Fw | CTCACGCCGACGAACAAAAAgagaaaaaatggctgcatcaagggc |
| pCA_ibpA_alpha43/63::B_ins_Rv | CAGGTTAGCACCACGAACATGAATGTTCTCagccagcgtaaagctcagg |
| pCA_ibpA_alpha43/63::B_vec_Fw | GTTCGTGGTGCTAACCTGGTAAATGGTTTGCTGTATATCGATCTCGAAC |
| pCA_ibpA_alpha43/63::B_vec_Rv | GTTCGTCGGCGTGAGCACCTTTCACCACCAGCAGATTATCCTGGG |
| pCA_ibpA_alpha64/84::B_ins_Fw | GAACGCAAATTCCAGTTAGCTgaaaatatggaagtctctggcgcaacc |
| pCA_ibpA_alpha64/84::B_ins_Rv | CGCTTCCGGAATCACacgaattaaatcaatatgcagtaaaccgtttacg |
| pCA_ibpA_alpha64/84::B_vec_Fw | gtGTGATTCCGGAAGCGAAAAAACCGCGCCGTATCGAAATCAACtaaGG |
| pCA_ibpA_alpha64/84::B_vec_Rv | cAGCTAACTGGAATTTGCGTTCAAAGTTGCGTTCAGCGATGCCCTGG |
| pCA_ibpA_alpha43/51::B_Fw | gagaaaaaatggctgcatcaagggcttGCTGAACGCAACTTTGAACGC |
| pCA_ibpA_alpha43/51::B_Rv | CaagcccttgatgcagccattttttctcTTTTTGTTCGTCGGCGTGAG |
| pCA_ibpA_alpha52/63::B_ins_Fw | CTATCTGTACCAGGGCATCatgaatcagccatttagcctg |
| pCA_ibpA_alpha52/63::B_ins_Rv | CCACGAACATGAATGTTCTCagccagcgtaaagctcag |
| pCA_ibpA_alpha52/63::B_vec_Fw | GAGAACATTCATGTTCGTGGTG |
| pCA_ibpA_alpha52/63::B_vec_Rv | GATGCCCTGGTACAGATAGG |
| pCA_ibpA_alpha52/56::B_Fw | CatgaatcagccatttGAACGCAAATTCCAGTTAGC |
| pCA_ibpA_alpha52/56::B_Rv | CaaatggctgattcatGATGCCCTGGTACAGATAGG |
| pCA_ibpA_alpha57/63::B_Fw | agcctgagctttacgctggctGAGAACATTCATGTTCGTGG |
| pCA_ibpA_alpha57/63::B_Rv | agccagcgtaaagctcaggctAAAGTTGCGTTCAGCGATG |
| pCA_ibpA_E92A_Fw | GGCATCGCTgcgCGCAAC |
| pCA_ibpA_E92A_Rv | GTTGCGgcgAGCGATGCC |

| | |
|---|---|
| pCA_ibpA_R93A_Fw | GGGCATCGCTGAAgcgAACTTTG |
| pCA_ibpA_R93A_Rv | CTGGAATTTGCGTTCAAAGTTcgcTTCAG |
| pCA_ibpA_N94A_Fw | GCTGAACGCgcgTTTGAACGC |
| pCA_ibpA_N94A_Rv | TGGAATTTGCGTTCAAAcgcGCG |
| pCA_ibpA_F95A_Fw | GCTGAACGCAACgcgGAACGC |
| pCA_ibpA_F95A_Rv | CTGGAATTTGCGTTCcgcGTTGCG |
| pCA_ibpA_R93K_Fw | GGGCATCGCTGAAaaaAACTTTGAACGC |
| pCA_ibpA_R93K_Rv | TTtttTTCAGCGATGCCCTGGTACAG |
| pCA_ibpA_R93N_Fw | GGCATCGCTGAAaacAACTTTGAACGC |
| pCA_ibpA_R93N_Rv | TTgttTTCAGCGATGCCCTGGTACAG |
| pCA_ibpA_R93W_Fw | GCATCGCTGAAtggAACTTTGAACGC |
| pCA_ibpA_R93W_Rv | TTccaTTCAGCGATGCCCTGGTACAG |
| pCA_ibpA_R93Y_Fw | CATCGCTGAAtatAACTTTGAACGCAAATTCCAG |
| pCA_ibpA_R93Y_Rv | TTataTTCAGCGATGCCCTGGTACAGATAGG |
| pCA_ibpA_R93F_Fw | GGCATCGCTGAAtttAACTTTGAACGCAAATTC |
| pCA_ibpA_R93F_Rv | TaaaTTCAGCGATGCCCTGGTACAGATAGG |
| pCA_ibpA_R93I_Fw | CTGTACCAGGGCATCGCTGAAattAACTTTG |
| pCA_ibpA_R93I_Rv | GTTCTCAGCTAACTGGAATTTGCGTTCAAAGTTaatTTC |
| pCA_ibpA_R93L_Fw | CCAGGGCATCGCTGAActgAACTTTG |
| pCA_ibpA_R93L_Rv | CAGCTAACTGGAATTTGCGTTCAAAGTTcagTTC |
| pCA_ibpA_R93V_Fw | CCAGGGCATCGCTGAAgtgAACTTTG |
| pCA_ibpA_R93V_Rv | CAGCTAACTGGAATTTGCGTTCAAAGTTcacTTC |
| pCA_ibpA_R93G_Fw | CCAGGGCATCGCTGAAggcAAC |
| pCA_ibpA_R93G_Rv | GAATTTGCGTTCAAAGTTgccTTCAGCG |
| pCA_ibpA_R93P_Fw | CCAGGGCATCGCTGAAccgAAC |
| pCA_ibpA_R93P_Rv | GGAATTTGCGTTCAAAGTTcggTTCAGC |
| pCA_ibpA_R93C_Fw | CAGGGCATCGCTGAAtgcAACTTTG |
| pCA_ibpA_R93C_Rv | GCTAACTGGAATTTGCGTTCAAAGTTgcaTTC |
| pCA_ibpA_R93M_Fw | ACCAGGGCATCGCTGAAatgAACTTTG |
| pCA_ibpA_R93M_Rv | CAGCTAACTGGAATTTGCGTTCAAAGTTcatTTC |
| pCA_ibpA_R93H_Fw | GTACCAGGGCATCGCTGAAcatAACTTTG |
| pCA_ibpA_R93H_Rv | CAGCTAACTGGAATTTGCGTTCAAAGTTatgTTC |
| pCA_ibpA_R93D_Fw | GTACCAGGGCATCGCTGAAgatAACTTTG |
| pCA_ibpA_R93D_Rv | CTCAGCTAACTGGAATTTGCGTTCAAAGTTatcTTC |
| pCA_ibpA_R93E_Fw | GAAgaaAACTTTGAACGCAAATTCCAGTTAGCTG |
| pCA_ibpA_R93E_Rv | CAAAGTTttcTTCAGCGATGCCCTGGTACAG |
| pCA_ibpA_R93Q_Fw | GAAcagAACTTTGAACGCAAATTCCAGTTAGCTG |
| pCA_ibpA_R93Q_Rv | CAAAGTTctgTTCAGCGATGCCCTGGTAC |
| pCA_ibpA_R93S_Fw | GAAagcAACTTTGAACGCAAATTCCAGTTAGC |
| pCA_ibpA_R93S_Rv | CAAAGTTgctTTCAGCGATGCCCTGG |
| pCA_ibpA_R93T_Fw | GAAaccAACTTTGAACGCAAATTCCAGTTAGC |
| pCA_ibpA_R93T_Rv | CAAAGTTggtTTCAGCGATGCCCTGGTAC |
| pBAD_sfGFP_Vec_Fw | ATGAGTAAAGGAGAAGAACTTTTCACTGGAGTTGTCC |

| | |
|---|---|
| pBAD_sfGFP_Vec_Rv | ATGGAGAAACAGTAGAGAGTTGCGATAAAAAGCGTCAG |
| pBAD_C.neteri ibpA 5' UTR_Fw | aggtaaccaaaacgagtacgccgattgg |
| pBAD_C.neteri_ibpA 5' UTR_Rv | gttttggttacctATGGAGAAACAGTAGAGAGTTGCGATAAAAAGCG |
| pBAD_V.harveyi_ibpA 5' UTR_Fw | AACTCTCTACTGTTTCTCCATcgctaaacgagatcgctc |
| pBAD_V.harveyi_ibpA 5'UTR_Rv | AAAGTTCTTCTCCTTTACTCATaatctatcctcttttaagcaatagtc |
| pCA_ibpA_C.neteri_R93A_Fw | AAgcgAACTTCGAGCGTAAATTTCAAC |
| pCA_ibpA_C.neteri_R93A_Rv | CGAAGTTcgcTTCCGCGATG |
| pCA_ibpA_V.harveyi_R94A_Fw | GAGgcgGATTTTGAACGTAAATTCCAAC |
| pCA_ibpA_V.harveyi_R94A_Rv | CAAAATCcgcCTCTGCAATGCC |
| pCA_ibpA_R97A_Fw | CGCTGAACGCAACTTTGAAgcgAAATTCC |
| pCA_ibpA_R97A_Rv | CGAACATGAATGTTCTCAGCTAACTGGAATTTcgcTTC |
| pCA_ibpA _R83A_Fw | GAGgcgACCTATCTGTACCAGGGC |
| pCA_ibpA_R83A_Rv | GGTcgcCTCTTTTTGTTCGTCGGC |
| pCA_ibpA _K98A_Fw | CGCgcgTTCCAGTTAGCTGAGAAC |
| pCA_ibpA_K98A_Rv | GAAcgcGCGTTCAAAGTTGCGTTCAG |
| pCA_ibpA_R97K_Fw | CGCAACTTTGAAaaaAAATTCCAGTTAGCTGAGAAC |
| pCA_ibpA_R97K_Rv | TTTtttTTCAAAGTTGCGTTCAGCGATGCC |
| pCA_ibpA_K98R_Fw | GAACGCcgcTTCCAGTTAGCTGAG |
| pCA_ibpA_K98R_Rv | gcgGCGTTCAAAGTTGCGTTCAG |
| pCA_ibpA_Y34R_Fw | GGCGGCcgtCCTCCGTATAACG |
| pCA_ibpA_Y34R_Rv | GGacgGCCGCCATTACTCTGGC |
| pCA_ibpA_Y34A_Fw | GGCGGCgctCCTCCGTATAACG |
| pCA_ibpA_Y34A_Rv | GGagcGCCGCCATTACTCTGGC |
| pCA_ibpA_Y34F_Fw | GCCAGAGTAATGGCGGCtttC |
| pCA_ibpA_Y34F_Rv | GTTCAACGTTATACGGAGGaaaGCCG |
| pCA_ibpA_Y34W_Fw | GCCAGAGTAATGGCGGCtggCC |
| pCA_ibpA_Y34W_Rv | GTTCAACGTTATACGGAGGccaGCCG |
| pCA_ibpA_Y34H_Fw | GAGCCAGAGTAATGGCGGCcatCC |
| pCA_ibpA_Y34H_Rv | GTTCAACGTTATACGGAGGatgGCCGC |
| pCA_ibpA_IEI-AEA_Fw | GTgccGAAgccAACtaaGGGTCGACCTGCAGCCAAGC |
| pCA_ibpA_IEI-AEA_Rv | ttaGTTggcTTCggcACGGCGCGGTTTTTTCGCTTC |
